# High-resolution HLA and MICA/MICB typing of the 1000 Genomes panel: a resource for precision medicine

**DOI:** 10.64898/2026.10.22.761181

**Authors:** Chul-Woo Pyo, Wyatt C. Nelson, Ruihan Wang, Yue Song, Alexander Thomas, Dayoung Kim, Eun Bin Go, Shalini Pereira, Akiko Ishitani, Daniel E. Geraghty

## Abstract

High-resolution characterization of human leukocyte antigen (HLA) and MICA/MICB variation is important for transplantation, immunogenetics, and the interpretation of disease associations within the major histocompatibility complex (MHC). However, publicly available population reference datasets remain limited in their resolution and coverage of these loci. We performed high-resolution typing of the 1000 Genomes Phase 3 panel across 13 HLA and MIC loci, generating HLA genotypes for 2,486 individuals, extended HLA-A, -B, and -C allele assignments, and MICA/MICB genotypes for 2,393 individuals representing 25 populations and five continental super-populations. Concordance with previously reported HLA genotypes was 96.2-97.4% at two-field resolution, and independent retyping confirmed the present assignments for samples exhibiting reciprocal genotype discrepancies with an earlier dataset. Analysis of extended class I variation within defined assay boundaries revealed that most of the 78 common three-field HLA lineages were dominated by a single extended sequence group. However, eleven lineages exhibited substantial heterogeneity, with the predominant group accounting for less than 80% of observed copies. The distinguishing sequence differences were noncoding, involving intronic and 3′ untranslated regions, and their distributions varied among ancestral populations. These findings identify recurrent, population-relevant variants for which functional consequences remain to be established. Population analyses demonstrated extensive allelic diversity at classical class I loci and DRB1, whereas the greatest between-population differentiation occurred at selected class II and DRB-linked loci. Most common alleles were shared among populations but differed substantially in frequency. Multilocus haplotypes revealed characteristic population-associated combinations of HLA and MIC alleles and provided the basis for predicting MIC genotypes from existing HLA typing. Using two-field HLA-B and HLA-C genotypes, five-fold cross-validation achieved exact genotype prediction accuracies of 77.3% for MICA and 53.6% for MICB, compared with 4.6% and 21.6%, respectively, without HLA information. Together, these results provide an extended-resolution HLA and MIC reference resource, demonstrate the limited routine utility of universal fourth-field typing while identifying specific variants for functional investigation, and establish the potential for recovering MIC variation from existing HLA-typed cohorts. The resource also supports future investigations of MHC variation underlying disease associations and transplantation outcomes.

## 1 Introduction

The practice of using genetic information from patients as guidance for clinical application, so-called precision medicine, has been used for over 40 years in hematopoietic cell transplantation (HCT) (1, 2), by examining donor and recipient for allelic differences in the human leukocyte antigen (HLA) gene system (3, 4). As the HLA class I and II loci are the most polymorphic loci in the human genome and play a central role in the adaptive immune response, the functional role that HLA allelic differences have as a direct effect on HCT outcomes has long been evident (5). More recent work has suggested other polymorphic gene systems affect HCT outcomes directly and indirectly through both allelic variation and immune response function including MICA/MICB (6, 7), KIR (8, 9), and genome-wide (10, 11). More recent emerging cellular immunotherapies that have in context evolved from HCT therapies are likely also affected by immune response gene allelic variations (12–15).

Outside of HCTs but within the framework of precision medicine is the goal of assigning genetic variation to common diseases through genome-wide association studies (16). Relevant to HLA and MICA/MICB, many disease-association signals map to chromosomal region 6p21.3, the major histocompatibility complex (MHC), where the HLA and MICA/MICB genes reside alongside numerous other genes involved in immune function (17). Indeed, targeted HLA and disease associations have been carried out continuously for decades (18–21), including early investigations of type 1 diabetes (T1D) more than 50 years ago (22–24). With these considerations, extending precision medicine into the mainstream of clinical practice requires critical elements in both directed novel technologies and the creation of population datasets that can inform the details of genetic variation at HLA and other immune response loci.

Given the role of HLA and MICA/MICB in the immune response, which in turn is central to most common diseases, we set out to build and analyze a panel of relevant genetic data based on publicly available resources. The 1000 Genomes Project was targeted as among the largest public catalogue of human variation and genotype data that is available for further work through the International Genome Sample Resource (IGSR), including DNA samples. We analyzed 2,486 individuals from 25 populations (25, 26). Several aims motivated this work: to test a generalizable next-generation typing protocol (27), to refine existing 1000 Genomes data resources for population studies and data mining studies, and as a supplement to existing HLA and immune response genetic resources (28). By generating high-resolution HLA typing, extended class I assignments within assay coverage, and exon-focused two-field typing for MICA and MICB it was possible to contribute three key extensions of the 1000 Genomes resources. First, a summary of the utility of 3-field and 4-field HLA class I allele groups and haplotypes extended across the 25 ancestral populations was generated toward relevance to basic and clinical applications. Intronic variation in HLA class I was examined with respect to its potential utility in the HCT clinical setting and in association analyses in the research setting. A second extension derived from aggregate comparisons of HLA and MICA/MICB types was made toward describing allele-frequency and haplotype differences among ancestral groups. Both individual allele types and HLA and MICA/MICB haplotypes were examined across individual populations and broader population groups. A third extension follows from the MIC data specifically. Published MIC imputation resources use genome SNP data (29); prediction from HLA types alone addresses a complementary setting. Because MICA lies within the class I region in strong linkage disequilibrium with HLA-B (30, 31), we used the paired HLA and MIC typings available for these samples to construct and cross-validate a predictor of MICA and MICB allele types from HLA-B and HLA-C type alone. This evaluates the accuracy attainable within this reference without additional MIC genotyping and, because HLA typing is already held by transplant cohorts and donor registries that lack MIC typing, defines the extent to which MIC can be recovered retrospectively from existing clinical datasets. These findings were discussed with relevance to personalized medicine that is represented by immunotherapies in general and HCT therapies in particular.

## 2 Materials and methods

### 2.1 Study cohort and analytical scope

The 1000 Genomes Project Phase 3 panel was used for HLA and MIC typing with the study workflow. The analytical cohort comprised 2,486 individuals representing 25 sub-populations assigned to five continental super-populations: African (AFR), Admixed American (AMR), East Asian (EAS), European (EUR), and South Asian (SAS). The final HLA dataset thus comprised 2,486 individuals, whereas MICA/MICB data were acquired for 2,394 individuals. Complete paired HLA-B/C and MIC genotypes were available for 2,393 individuals, who were used for MIC prediction and the paired MIC frequency summaries. Allele calls were obtained at all eight non-paralogous HLA loci for all 2,486 individuals, and at DRB3, DRB4 and DRB5 for all individuals carrying the corresponding paralogue, and at MICA and MICB for 2,393 individuals. Denominators nevertheless differ across loci and gene systems: at DRB3, DRB4 and DRB5 the denominator is the number of observed paralogue copies rather than the full cohort, since these genes are present only on defined DRB1 haplotypes, which reflects genomic organization rather than incomplete typing. All downstream allele-frequency, diversity, haplotype and MIC prediction analyses were therefore conducted using locus-specific denominators rather than assuming a single uniform sample size across all genes. The study design and population framework were based on the 1000 Genomes Project Phase 3 resource (25).

### 2.2 HLA and MICA/MICB typing

HLA and MICA/MICB genotyping was performed using the ScisGo-HLA-v6 and ScisGo-MICAB-v3 typing kits (Scisco Genetics, Inc., Seattle, WA). These kits provided universal targeted amplicon workflows designed to generate exon-level sequence evidence across all targeted loci within an integrated laboratory and bioinformatics framework (Supplementary Figure 1). Sequencing data were analyzed using IPD-IMGT/HLA release 3.66.0 (October 2026) for high-resolution allele assignment. The resulting genotypes were used for all analyses reported in this study. The HLA and MICA/MICB technologies follow an essentially identical two-stage PCR strategy consisting of four coordinated multiplex reactions. The protocols started with stage 1 assembly of 2 µL DNA and 8 µL S1 buffer addition to HLA or MICA/MICB reaction plates followed by stage 1 PCR according to the ScisGo protocol-defined PCR cycling program. At completion, plates are removed and 10 µL of S2 buffer is added to each well followed by stage 2 PCR, finishing with pooling and buffer exchange. Following amplification, products from the four reactions were combined within reaction sets, column-purified, quantified, normalized, and prepared for paired-end Illumina sequencing on a MiSeq or equivalent platform (Illumina, San Diego, CA).

Sequence data were processed using the centralized cloud-based bioinformatics environment Sciscloud.net (Scisco Genetics, Inc., Seattle, WA). Allele names and extended candidate lists were reported against IPD-IMGT/HLA release 3.66.0 (October 2026) (32). The analytical workflow standardized read preprocessing, reference alignment or allele matching, exon-level evidence evaluation, quality control, allele assignment, and report generation. Centralized execution ensured consistent application of reference resources, quality-control thresholds, interpretation logic, and reporting procedures across sequencing runs.

Final allele calls were reviewed in a desktop or web-based application, GeMS-UI (Scisco Genetics, Inc. Seattle WA) that enabled inspection of supporting read evidence, targeted editing of provisional assignments when necessary, which occurred at an average of three samples per 96-sample plate, and export of curated report-ready outputs. The standard kit configuration provides exon-focused evidence with additional intronic and partial terminal-region coverage, but does not establish every named fourth-field distinction. The genomic regions interrogated by the standard ScisGo-HLA-v6 assay encompass all currently known HLA fourth-field sequence variants that have been demonstrated to affect gene expression or protein function, as documented in the ScisGo-HLA-v6 kit supporting documentation (Scisco Genetics, Inc., Seattle, WA). Consequently, although the standard assay does not distinguish every fourth-field allele catalogued in IPD-IMGT/HLA, its sequence coverage includes the established functionally consequential variants. The supplementary reactions described below were designed to extend sequence coverage and characterize additional noncoding variation, rather than to recover previously established functional distinctions absent from the standard assay.

This design supported integrated high-resolution interpretation; throughput and cost were not benchmarked against long-range PCR approaches (Supplementary Figure 1). Each distinct lot and shipment of kits was tested using a control DNA panel, including study controls and available reference standards. Positive and negative controls were placed on each 96-sample plate to verify orientation and plate identity, and duplicate study samples were distributed across plates. Control and duplicate genotypes were reviewed as part of the laboratory quality-control procedure.

The loci examined included the class I genes HLA-A, HLA-B and HLA-C; the class II genes DPA1, DPB1, DQA1, DQB1 and DRB1; the DRB paralogues DRB3, DRB4 and DRB5; and MICA and MICB, yielding 13 analytical loci in total. The exported labels include the intended high-resolution assignments and some shorter or provisional labels, which were retained as reported.

### 2.3 Extended HLA-A, HLA-B, and HLA-C four-field typing and phasing

To enable extended 4-field resolution for HLA-A, HLA-B and HLA-C, two supplementary multiplex reactions were added to the standard four-reaction ScisGo workflow. These reactions targeted intron 3, which the standard kit configuration does not fully cover, together with the remaining intronic regions and partial 5′ and 3′ untranslated regions, extending intronic and terminal-region sequence evidence for the samples reported here. Coverage remained insufficient to distinguish some named HLA-B and HLA-C candidates.

Extended 4-field typing was performed by integrating sequence data from the standard exon-focused assay with reads generated from the supplementary intron-targeting reactions. After preliminary allele assignment, reads supporting the candidate HLA-A, HLA-B, and HLA-C alleles were collected on a locus-by-locus basis and assembled using data from both the standard typing reactions and the supplementary intronic reactions. Consensus sequences were then generated across the extended class I gene regions.

The resulting consensus sequences were compared against IPD-IMGT/HLA release 3.66.0 (October 2026) reference alleles to identify the best-supported extended assignments. This strategy enabled exon-based typing evidence from the standard workflow to be combined with additional intronic and untranslated-region sequence information, thereby improving phasing across the extended gene structure and supporting high-resolution HLA-A, HLA-B, and HLA-C allele assignment (32, 33). Ambiguities were retained when weak or missing read coverage at candidate-discriminating positions left the extended assignment unresolved. Extended calls were summarized as sequence groups supported within the typing interval; differences outside that interval did not define additional observed groups. The detailed comparison and grouping procedures are described in Supplementary Methods.

### 2.4 Allele frequencies, diversity, and population differentiation

Typing results were represented as one record per sample, locus and observed copy. HLA analyses retained the exported canonical allele labels, including shorter and longer reported forms. MICA and MICB types used up to two reported numeric fields, retaining expression suffixes. In the supplied MIC exports, third-field extensions represent intronic variation; these extensions were retained separately as source annotations and were not counted as additional primary MIC types. Shorter labels and provisional tags within the retained fields were preserved without inferring missing fields. Common alleles were defined by a pooled global frequency of at least 1% at the relevant locus.

Expected heterozygosity was used to summarize within-group diversity because it reflects both the number of observed alleles and the evenness of their frequencies, rather than richness alone. It describes the probability that two independently sampled allele copies have different labels (34). Values were calculated from all observed allele labels within each super-population, then summarized by an unweighted mean across the five groups so that larger groups did not dominate the locus comparison. For DRB paralogues, the calculation was conditional on observed copies and did not treat structural absence as a conventional allele.

Pairwise FST was used to distinguish within-group diversity from differentiation between groups. This comparison asks whether allele copies sampled from different super-populations are more dissimilar than copies sampled within either group. A Hudson-type allele-frequency estimator incorporated finite-sample corrections to within-group diversity and used all observed alleles (35). Negative estimates were retained rather than truncated, and locus summaries gave equal weight to the ten super-population pairs. These estimates describe allele-frequency differentiation; they do not independently identify demographic or selective causes.

Allele-specific association with super-population was summarized by Cramér’s V from a table comparing the counts of each allele and all other alleles across the five groups. V was chosen to rank the magnitude of association on a standardized scale, rather than rank significance levels that also depend on sample size (36). Classification was hierarchical: alleles observed in only one group were population-private; among the remainder, V of at least 0.30 defined high differentiation, V from 0.15 to less than 0.30 defined moderate differentiation, and V below 0.15 with presence in all five groups defined the public class. Remaining alleles present in four groups were broadly shared; those present in two or three groups formed the residual super-population-enriched class. Enrichment ratios supplemented V by expressing the highest group frequency relative to the unweighted mean in the other four groups. They indicate the direction and relative size of enrichment, rather than provide an additional significance test. These categories describe the sampled panel and do not establish selection or universal geographic restriction.

### 2.5 Data preprocessing and allele normalization

Data integration and normalization were conducted within a standardized preprocessing framework. HLA and MICA/MICB typing results were transformed into a unified long-format table, with one record per sample, locus, and recorded allele copy. For allele-frequency, diversity, differentiation and association analyses, the Allele field retained the exported HLA labels and the MIC types at up to two reported fields. Original MIC labels, intronic extensions and provisional-tag flags were retained in separate copy-level and label-level inventories in the released source-annotation data. Fourth-field information was retained separately only for the HLA-A, HLA-B, and HLA-C predictability analysis. Each sample record was merged with the corresponding population and super-population annotations, and DRB345-linked calls were disaggregated into locus-specific DRB3, DRB4, and DRB5 records for downstream locus-level analyses. Except where noted, all analyses were conducted in Python. Data processing and table generation used pandas and NumPy; figures were produced with Matplotlib. Reported allele assignments were retained as generated using IPD-IMGT/HLA release 3.66.0, without subsequent nomenclature substitution.

Allele assignments were reviewed using the available variant information and sequence-coverage annotations before analysis. All analyses used the curated study calls. Population and super-population metadata were linked by sample identifier. Variant and coverage annotations are provided in Supplementary Table 7.

### 2.6 Multilocus haplotypes and extended resolution summaries

We estimated and summarized expectation-maximization (EM) haplotypes for six blocks: class I (A∼B∼C), the MIC-linked block (B∼C∼MICA∼MICB), class II core (DRB1∼DQA1∼DQB1∼DRB345), extended class II (DPA1∼DPB1∼DQA1∼DQB1∼DRB1∼DRB345), full HLA (nine loci including DRB345) and combined HLA-MIC (eleven loci). The first three are reported in the main figure; the extended class II, full HLA and combined HLA-MIC blocks are reported as supplementary because their higher diversity and lower retained frequency mass limit interpretation. DRB345 is a synthetic locus representing an exported DRB3, DRB4 or DRB5 call or an absence state. EM was used because the genotypes were unphased: the two recorded copies at each locus could not be assigned directly to the same chromosome across loci. The procedure considered compatible haplotype pairs within each super-population and iteratively updated their frequencies, retaining phase uncertainty rather than treating copy-column order as known phase (37). Samples had to have both recorded copies at every locus required by a block. The source estimation procedure pooled rare alleles below 0.5% separately within copy columns and exported haplotypes at frequencies of at least 0.1% after excluding pooled-rare haplotypes. For MIC-containing blocks, retained estimates with identical two-field MIC types were summed within each super-population, and conditional diversity was recalculated after this aggregation without re-estimating phase.

Because the exported frequencies do not sum to one, diversity was calculated after rescaling the retained estimates to a complete conditional distribution. Effective haplotype number expresses diversity as the number of equally frequent haplotypes that would give the same probability of sampling a matching pair, and therefore emphasizes the concentration of common combinations (38); Shannon entropy, reported in bits, summarizes uncertainty across the frequency distribution and gives relatively more weight to less common combinations (39). Retained frequency mass and sample size were reported alongside them because neither measure recovers haplotypes omitted from the export. Heatmaps show eight leading retained haplotypes chosen from all five groups’ leading haplotypes and then by the unweighted mean group frequency; displayed frequencies retain the original EM scale.

The class I resolution analysis included 24 common HLA-A, 30 HLA-B and 24 HLA-C lineages with pooled frequency ≥1% in the 2,486-individual panel. The extended HLA-A, -B and -C typing intervals covered coding regions, introns and proximal genomic flanks, extending 300, 284 and 319 bp before the ATG start codon and 318, 586 and 394 bp after the stop codon, respectively. To avoid counting differences outside assay coverage, reported fourth-field candidates at each locus were matched to IPD-IMGT/HLA 3.66.0 genomic references and compared within these intervals. Complete within-interval references with identical sequences defined one sequence group; known nucleotide or insertion/deletion differences within the interval separated groups. Differences found only outside the interval did not create separate observed groups. Shorter references were compared at available positions without filling unknown bases, as detailed in Supplementary Methods. These groups summarize the available sequence evidence within the assay, rather than establishing every full-genomic fourth-field distinction.

Each reported allele copy was mapped to the group or groups represented by its ambiguity candidates. A copy was counted once when its candidates supported a single group, or when an explicit sequencing-confirmed group assignment was available; the number of candidate labels did not increase its count. Copies without a single supported group retained their reported calls and annotations but did not enter group-fraction denominators. Grouping did not replace the primary HLA types used for allele-frequency or haplotype analyses. Each group was displayed using the numerically lowest complete-reference member, or the lowest shorter-reference member if none was complete. A dagger (†) marks the representative of a sequence group containing two or more distinct listed allele labels; single-member groups are unmarked. The symbol is not part of HLA nomenclature. Supplementary Table 1 lists group members, assignment status and denominators. The common-lineage threshold of pooled frequency ≥1% required at least 50 of 4,972 primary allele copies per locus, excluding singleton three-field lineages from this analysis. Within an eligible lineage, every assigned sequence group was retained, including groups observed once. For each common lineage, group counts were summed in the pooled cohort and separately within each super-population. The dominant-group fraction was the count of the largest group divided by the number of copies assigned to any group for that lineage. Lineages were classified as predictable at fractions of at least 95%, mostly predictable at 80% to less than 95%, and uncertain below 80%. Mean fractions were unweighted across common lineages. The same group definitions were used for pooled and super-population summaries, so ancestry comparisons reflected group frequencies rather than a redefinition of groups. Figures 1–2 and Tables 1A–1B summarize these fractions and group compositions; Supplementary Table 1 provides the counts and denominators. Cells with fewer than five assigned copies were masked in population figures. The eleven globally uncertain lineages were examined by their sequence-group compositions. Differences in sequence-group distributions between super-populations were summarized by mean pairwise Jensen-Shannon divergence in bits for super-populations with at least three assigned copies. This symmetric measure compares full sequence-group distributions, including differences beyond the leading group, and accommodates groups absent from one super-population (40). It was used as a descriptive complement to the dominant-group fraction, rather than as a significance test. Predictability refers to observed sequence groups within assay coverage and does not validate every full-reference fourth-field distinction.

**Figure 1.**
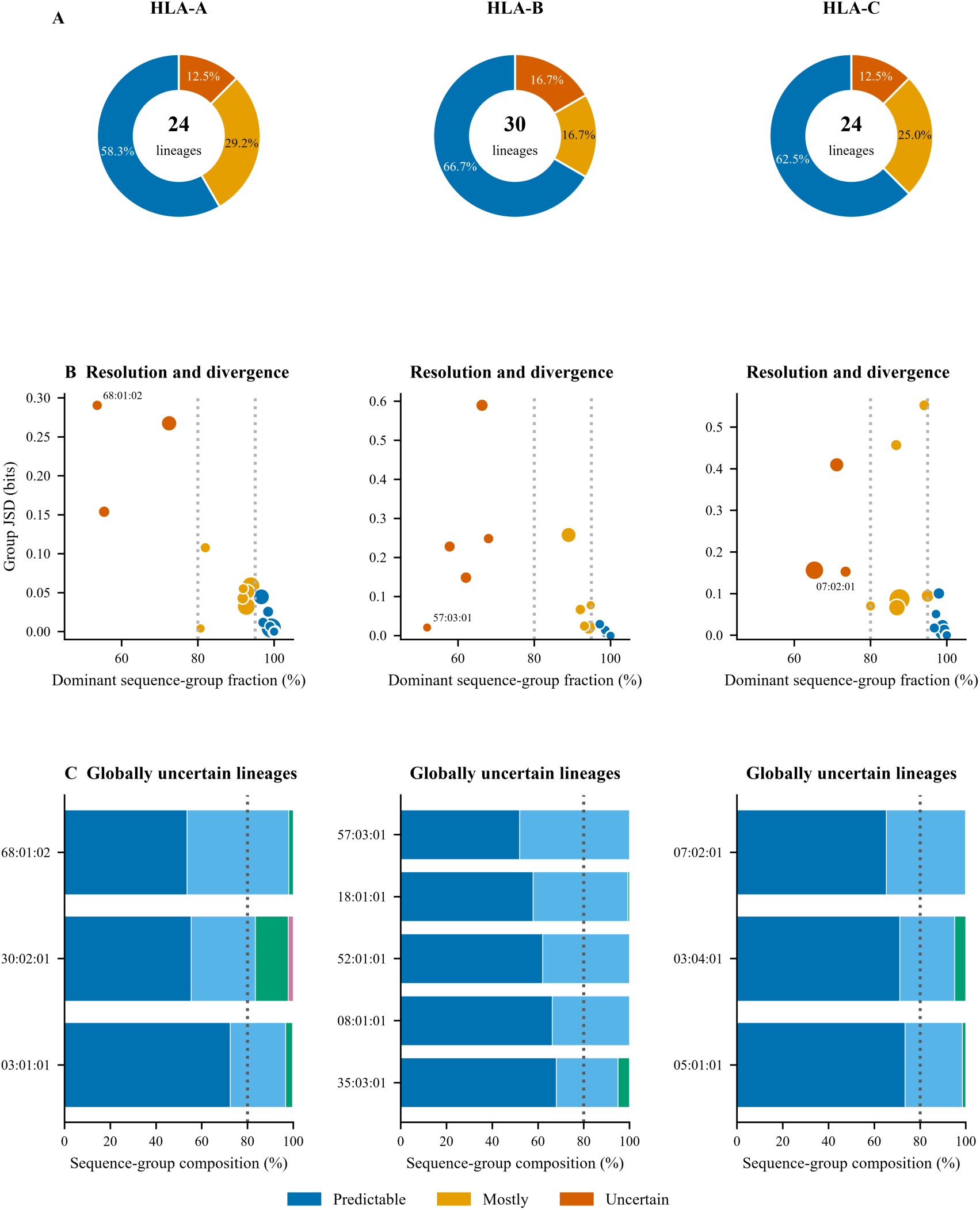
Three field to four field predictability within assay coverage. (A) Predictability classes for 24 common HLA-A, 30 HLA-B and 24 HLA-C lineages: predictable (dominant-group fraction ≥95%), mostly predictable (80% to <95%) and uncertain (<80%). (B) Dominant fraction versus mean pairwise Jensen-Shannon divergence between super-population sequence-group distributions. Bubble area increases with pooled lineage frequency. Dotted lines mark 80% and 95%. (C) Sequence-group composition of the eleven globally uncertain lineages, ordered by pooled frequency. Group memberships and assignment rules are supplied in Supplementary Table 1 and Supplementary Methods. Counts and means are in Table 1A, and the two leading representatives and their sequence differences are in Tables 1B–1C.

**Figure 2.**
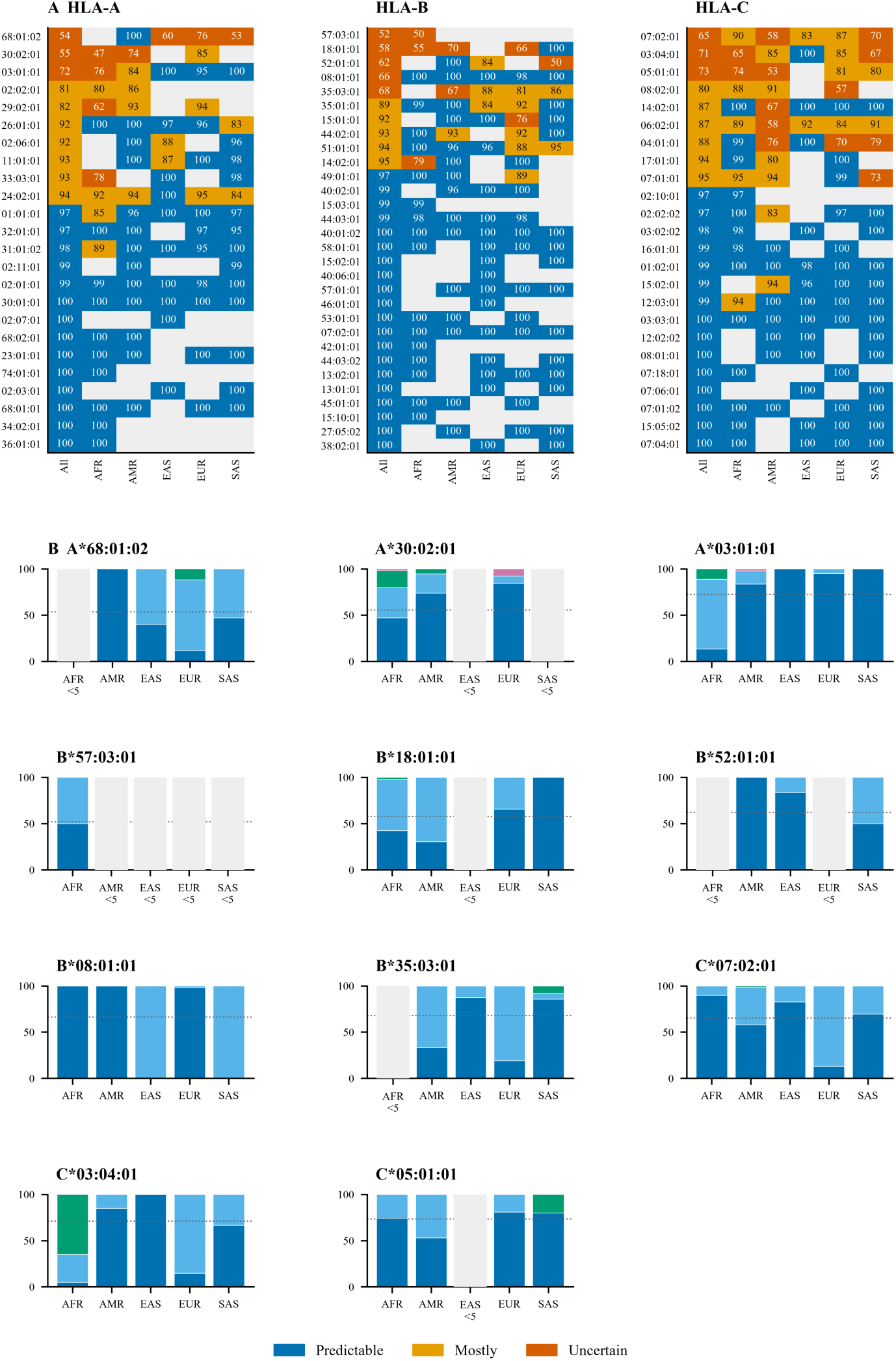
Ancestry dependence of extended class I resolution. (A) Dominant-group fractions (%) globally (All) and within each super-population. Colors use the Figure 1 thresholds; pale grey indicates fewer than five assigned copies. Fractions are rounded to integers for display, while classes use unrounded values. (B) Sequence-group composition of each globally uncertain lineage across the five super-populations. Colors are consistent within each lineage and with Figure 1C. Dotted lines show the pooled dominant fraction. Sparse groups are masked. Complete memberships and denominators are supplied in Supplementary Table 1; the grouping procedure is described in Supplementary Methods.

**Table 1A.**
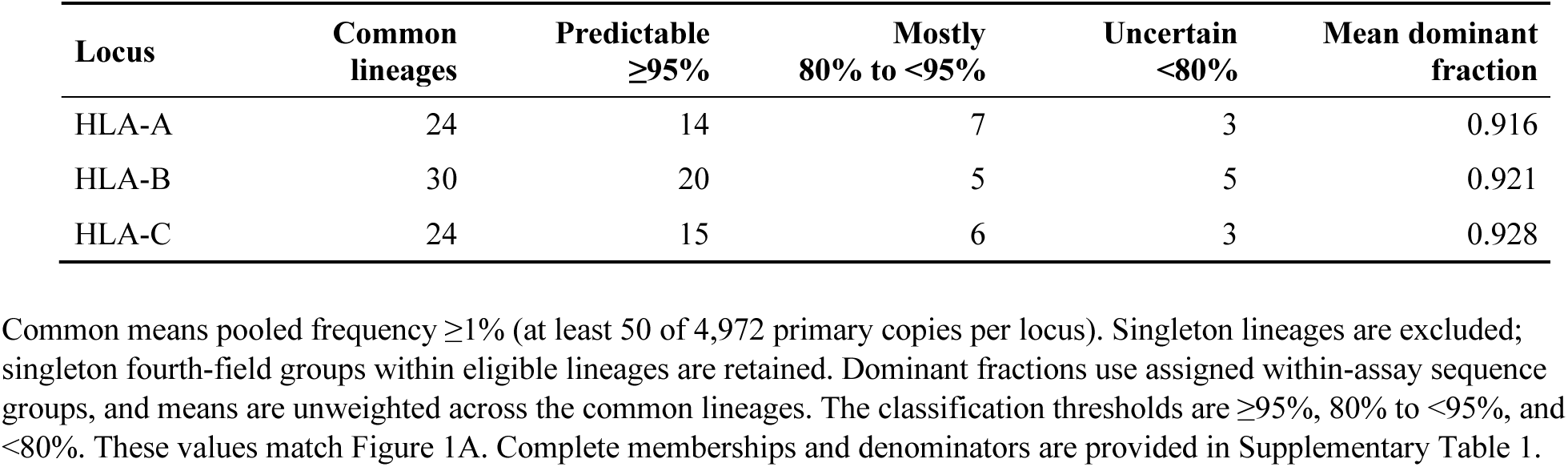
Class I resolution summary.

**Table 1B.**
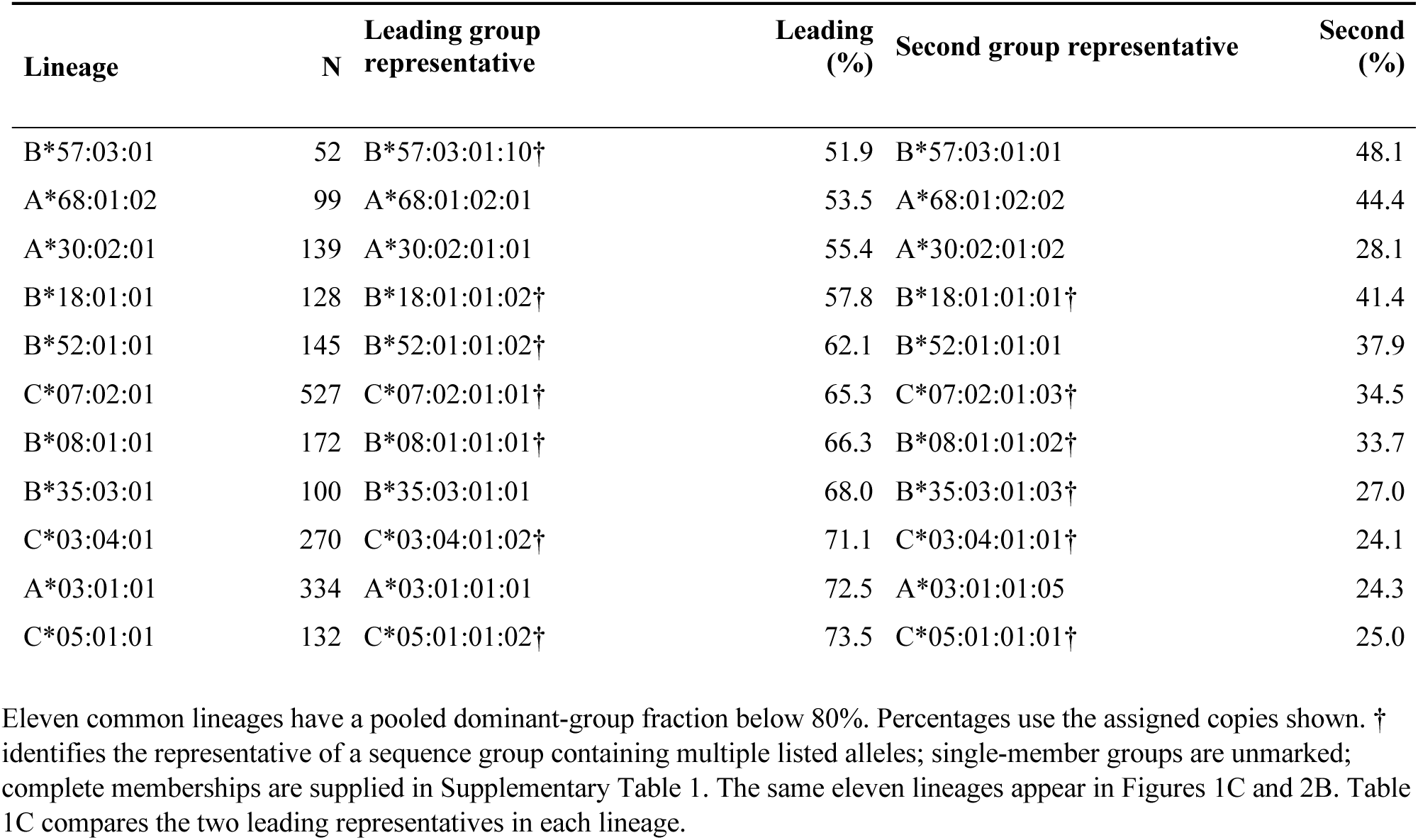
Uncertain common class I lineages.

To measure additional information across the cohort, rather than giving every lineage equal weight, we counted assigned allele copies outside the most frequent sequence group for each reported lineage. The primary baseline used pooled modal groups; a second used modal groups estimated separately within each super-population. Modal ties selected the numerically lowest representative. This separate cohort-weighted analysis included all reported lineages, including singletons. Primary lineage labels were retained without completing missing fields, and analyses restricted to common lineages or explicitly three-field labels were included as sensitivity checks. We then counted individuals with at least one known non-dominant group directly across their observed HLA-A, -B and -C copies, without assuming independence among loci or copies. Individuals without a known non-dominant copy but with an unresolved group remained indeterminate. Lower and upper bounds allowed these remaining indeterminate individuals to have no event or an event, respectively. These are descriptive, same-cohort measures of additional within-assay discrimination; they do not establish prospective prediction accuracy, clinical benefit or cost effectiveness (Supplementary Figure 6; Supplementary Table 1).

For each of the eleven uncertain lineages, the two most frequent pooled sequence groups were selected, and their four-field reference representatives were compared using IPD-IMGT/HLA 3.66.0 genomic alignments (Table 1C). Known differences within the typing interval were mapped to exons, introns and untranslated regions using canonical and allele-specific feature annotations. Table 1C reports the selected representatives and mapped differences; complete contrast records and coordinate definitions are provided in Supplementary Table 1 and Supplementary Methods. Sequence location was not treated as evidence of a functional effect.

**Table 1C.** Sequence differences between the two leading fourth-field groups.

| Three-field lineage | Leading and second four-field representatives | Within-interval differences (canonical positions) | Mapped region | Feature-relative positions |
| --- | --- | --- | --- | --- |
| HLA-A*68:01:02 | A*68:01:02:01<br>A*68:01:02:02 | 3070 G→A | Intron 7 | Intron 7: 41 |
| HLA-A*30:02:01 | A*30:02:01:01<br>A*30:02:01:02 | 3070 A→G | Intron 7 | Intron 7: 41 |
| HLA-A*03:01:01 | A*03:01:01:01<br>A*03:01:01:05 | 2906 C→T | Intron 6 | Intron 6: 67 |
| HLA-B*57:03:01 | B*57:03:01:10†<br>B*57:03:01:01 | 2530 C→T; 3314 T→C | Intron 5 + 3' UTR | Intron 5: 194; 3' UTR: 354 |
| HLA-B*18:01:01 | B*18:01:01:02†<br>B*18:01:01:01† | 1410 C→T; 2454 G→A; 3298 C→T | Intron 3, 5 + 3' UTR | Intron 3: 134; Intron 5: 118; 3' UTR: 338 |
| HLA-B*52:01:01 | B*52:01:01:02†<br>B*52:01:01:01 | 3262 C→T | 3' UTR | 3' UTR: 302 |
| HLA-B*08:01:01 | B*08:01:01:01†<br>B*08:01:01:02† | 3087 T→C | 3' UTR | 3' UTR: 127 |
| HLA-B*35:03:01 | B*35:03:01:01<br>B*35:03:01:03† | 2991 C→T; 11-bp indel (2996–3006) | 3' UTR | 3' UTR: 31; 3' UTR: 36–46 (indel) |
| HLA-C*07:02:01 | C*07:02:01:01†<br>C*07:02:01:03† | 1963 G→C | Intron 3 | Intron 3: 473 |
| HLA-C*03:04:01 | C*03:04:01:02†<br>C*03:04:01:01† | 3528 C→G | 3' UTR | 3' UTR: 138 |
| HLA-C*05:01:01 | C*05:01:01:02†<br>C*05:01:01:01† | 1632 T→C | Intron 3 | Intron 3: 142 |

† marks representatives of groups containing multiple listed alleles; single-member groups are unmarked. The symbol is not part of an allele name. The two representatives are the leading and second most frequent groups in Table 1B. Positions are one-based in the canonical genomic records HLA-A\*01:01:01:01, HLA-B\*07:02:01:01 and HLA-C\*01:02:01:01, rather than ATG-relative numbering. Base changes run from the first representative to the second; indel coordinates describe the aligned tract. Only known differences within the typing interval are shown. Full records and feature annotations are provided in Supplementary Table 1. Sequence location does not establish a functional effect.

### 2.7 Prediction of MICA and MICB from HLA type

We tested whether HLA-B and HLA-C types could predict MICA and MICB genotypes in 2,393 individuals with complete paired calls. The primary analysis used two-field HLA-B/C input and pooled haplotype frequencies. Joint HLA-B∼HLA-C∼MICA∼MICB haplotype frequencies were estimated by expectation-maximization, which accounts for unknown chromosome phase (37). For each HLA genotype, we predicted the most likely pair of MICA alleles and, separately, MICB alleles. Targets used up to two reported MIC fields, excluding third-field intronic extensions while retaining expression suffixes.

Performance was assessed by five-fold cross-validation, with individuals distributed within sub-populations. Haplotype frequencies were re-estimated using training individuals only, and the held-out MIC calls were used solely to assess accuracy. A prediction was correct only when both MIC alleles matched, irrespective of order. Exact-genotype accuracy was therefore a binomial proportion over held-out individuals, and its two-sided 95% interval was calculated by the Wilson score method, which remains well behaved for proportions near 0 and 1 (41). Accuracy was compared with always predicting the most frequent MIC genotype in each training fold, without using HLA input. This majority-genotype baseline tests whether HLA typing provides information beyond MIC genotype frequencies alone. We also evaluated HLA-B alone to measure the added information from HLA-C and reported HLA-B/C performance by super-population. All rare-allele and unsupported-input predictions remained in the accuracy denominator. Detailed model settings, fallback rules, additional configurations, allele-level results and confidence analyses are provided in the Supplementary Methods and Supplementary Table 4.

### 2.8 Comparison with previous 1000 Genomes typing

We compared the current workbook allele pairs with the 2014 IGSR release accompanying Gourraud et al. (42) and final Supplementary Table 2 of Abi-Rached et al. (43). Records were joined by the exact public sample identifier and locus. Review codes replace individual identifiers in manuscript displays. HLA-A, -B, -C, -DRB1 and -DQB1 were compared at the first two explicitly written allele fields, retaining expression suffixes and ignoring the order of the two copies. Complete source labels were preserved.

Explicit matches and ambiguity-compatible matches were counted separately. A listed ambiguity matched when a complete pair of written alternatives was compatible with the other genotype, preserving both allele copies and homozygous multiplicity. A shared-prefix abbreviation such as 02:01/30 was read as the written alternatives 02:01 and 02:30. Compatible alternatives were not selected as resolved calls. Match percentages combined explicit and ambiguity-compatible matches among all compared individual-locus records. Missing, broad or malformed calls and repeated source rows were excluded; repeated rows were not merged or selected. Source annotation asterisks were retained in the raw display but were not allele fields. No genotype imputation, G/P grouping, historical-name substitution or sample reassignment was used. Supplementary Table 5 summarizes concordance and lists exclusions; Supplementary Table 6 supplies all nonmatching records and reciprocal profiles. Full source labels and compatible written pairs are retained in the released comparison data. Explicit-only and complete-label equality counts remain available separately. For records with explicit types at all five loci, the current and 2018 profiles were also compared across identifiers. The study dataset provided 12,430 genotype pairs across the five compared loci. To investigate reciprocal cross-individual genotype matches identified between the present study and the 2018 dataset, the affected DNA samples were independently re-obtained from the original sample repository and subjected to repeat HLA genotyping using the study typing workflow. Repeat allele assignments were compared with the genotypes reported here and the corresponding previously published records to assess the reproducibility of the sample-to-genotype assignments.

## 3 Results

### 3.1 Comparison with earlier typing of shared individuals

The current cohort shares 971 individuals with the 2014 release and 2,409 with the final 2018 release (Table 4). At two-field resolution, combined explicit and ambiguity-compatible match rates with the 2018 types were 97.1% for HLA-A, 96.4% for HLA-B, 97.2% for HLA-C, 97.4% for DRB1 and 96.2% for DQB1. Listed alternatives contributed 31, 15, 9, 3 and 28 matches, respectively. Table 4 distinguishes these from explicit matches rather than treating the ambiguous genotypes as resolved.

The 2014 release contained more ambiguous records. Including matches to written alternatives allowed comparison of 960–962 individuals per locus, with match rates of 91.8–95.4%. Broad or malformed labels, missing calls and repeated rows remained excluded (Supplementary Table 5). Among the 943 individuals shared by all three datasets, comparison of the two earlier releases yielded 96.1–99.1% combined match rates (Supplementary Table 5).

Twenty pairs showed reciprocal exact matches at all five explicitly reported two-field loci between the current and 2018 files: each current profile matched the 2018 profile written under the other member’s identifier. Full source records and recurring one-allele differences are supplied without assigning a cause, changing identifiers or imputing calls. To investigate these reciprocal discrepancies, 20 affected individuals were independently reacquired from the original DNA repository and retyped at all loci. The repeat typing reproduced the original genotype assignments for all 20 individuals, confirming the correspondence between the independently obtained samples and the genotypes reported in the present study. These results support the accuracy of the present sample assignments and suggest that the reciprocal discrepancies reflect errors in the earlier dataset, although their origin could not be determined.

### 3.2 Three-field to four-field predictability and covered sequence-group structure

High-resolution HLA typing is reported at either 3-field or 4-field resolution, the fourth field denoting polymorphism outside the coding sequence (44, 45). Because a single 3-field designation can correspond to multiple 4-field entries in the IPD-IMGT/HLA database, we asked how much of that catalogued subtype structure is realized in a population sample, and how reliably a 3-field designation predicts it. The analysis distinguishes sequence groups supported by the available reference positions from unresolved differences outside the typing interval.

Three-field and extended class I assignments were determined for the 25-population panel and assessed at three levels: the individual populations, the five super-populations (AFR, AMR, EAS, EUR, SAS), and the aggregate sample. The processed data contained 344 canonical class I labels, including shorter forms, and 427 observed assigned sequence groups. Splits were markedly uneven across lineages, with most common lineages accounted for by a single dominant group. Full group membership and assignment summaries are provided in Supplementary Table 1.

For comparison, IPD-IMGT/HLA release 3.66.0 catalogued 5,408 fourth-field alleles for HLA-A (1,538), HLA-B (1,989) and HLA-C (1,881), distributed across 410 three-field lineages, a mean of 13.2 catalogued subtypes per lineage. The present study resolved its 344 canonical class I labels into 427 assigned sequence groups within the defined typing intervals, a mean of 1.24. Catalogued fourth-field depth is also highly skewed: the median lineage carries two catalogued subtypes while A*02:01:01 alone carries 279. Most named class I alleles have never been resolved to four fields at all, so catalogued depth reflects the distribution of sequencing effort as well as the diversity present in sampled populations.

Because the number of fourth-field distinctions can depend on the extent of noncoding sequence represented in individual reference records, the analysis was restricted to sequence differences within explicitly defined genomic intervals covered by the typing assays. Alleles differing only outside these intervals were assigned to the same sequence group, preventing additional distinctions from being counted solely because more extensive reference sequence was available. The reported sequence groups therefore describe variation distinguishable within a standardized experimental interval.

For the main predictability analysis, we restricted attention to common lineages with a pooled frequency of at least 1%. This yielded 78 lineages across HLA-A, HLA-B and HLA-C. Group representatives identify the sequence groups observed within assay coverage, without implying that every full-genomic fourth-field distinction was resolved.

Restricting to common alleles, the analysis included 24 HLA-A, 30 HLA-B and 24 HLA-C common class I lineages (Figure 1; Table 1A). Most were highly informative for their dominant 4-field subtype. At HLA-A altogether, 14 were strongly predictive at the ≥95% threshold, seven mostly predictive at 80% to <95%, and three uncertain below 80%. At HLA-B, the corresponding counts were 20, five and five; at HLA-C, 15, six and three.

Mean dominant-subtype fractions were high at all three loci — 0.916 for HLA-A, 0.921 for HLA-B and 0.928 for HLA-C (Figure 1; Table 1A) — indicating that for most common class I alleles a 3-field designation captured the dominant 4-field subtype with high fidelity. HLA-A*02:01:01 illustrates the extreme case: of 650 assigned copies, 645 belonged to the group represented by HLA-A*02:01:01:01 and five to four other groups, giving 99.2% dominance. Complete assignment strings, group memberships and denominators are provided in Supplementary Table 1.

Cohort-level counting places these per-lineage fractions in context. Across all reported lineages, 1,206 of 14,716 assigned copies (8.2%) lay outside their pooled modal group, and at least one such copy was present in 981 of 2,486 individuals (39.5%, rising to 44.4% if every unresolved call is counted as a distinction). Because each individual contributes two copies at each of three loci, this individual-level proportion is close to the value expected from the per-copy rate alone. Restricting the analysis to the 78 common lineages gave 9.1% of copies and 37.9% of individuals. The distinctions recorded were concentrated in a small number of recurrent splits rather than distributed across the catalogued fourth-field range; per-locus rates, complete denominators and sensitivity analyses are supplied in Supplementary Figure 6 and Supplementary Table 1.

Using pooled modal groups, the observed individual proportion varied from 22.8% in EAS to 55.4% in EUR. Using ancestry-specific modal groups reduced the overall non-dominant copy fraction to 5.9% and the known individual proportion to 29.8% under the stated tie convention. Both baselines were estimated in this cohort. Modal-tie alternatives, unresolved-call bounds and complete denominators are provided in Supplementary Table 1; these frequencies measure discrimination within the observed resource, rather than independently validated imputation or a functional effect.

A minority of common alleles nonetheless concealed sequence-group heterogeneity behind a shared 3-field designation. The least predictable common lineage at each class I locus was HLA-B*57:03:01, whose leading subtype accounted for 51.9% of chromosomes, HLA-A*68:01:02 at 53.5%, and HLA-C*07:02:01 at 65.3% (Figure 1; Table 1B). Eleven lineages fell below the 80% threshold: HLA-A*68:01:02, HLA-A*30:02:01, HLA-A*03:01:01, HLA-B*57:03:01, HLA-B*18:01:01, HLA-B*52:01:01, HLA-B*08:01:01, HLA-B*35:03:01, HLA-C*07:02:01, HLA-C*03:04:01 and HLA-C*05:01:01 (Figure 1; Tables 1B–1C). In most of these the two leading groups occurred at comparable frequencies, so the uncertainty reflects a recurrent split rather than rare-group noise. The two leading four-field representatives in each lineage are shown in Table 1C. Their known differences within the typing interval were entirely noncoding: five contrasts involved introns alone, four involved the 3′ UTR alone, and two involved both. These mapped differences describe the selected reference sequences and do not establish functional effects.

The most consequential feature of this uncertain set was that uncertainty was not a fixed property of the lineage but depended on super-population background (Figure 2). Several globally uncertain lineages became more readily predictable within some super-populations while remaining split in others. Among groups with at least five assigned copies, six lineages reached at least 95% dominance in one or more super-populations: HLA-A*03:01:01, HLA-A*68:01:02, HLA-B*08:01:01, HLA-B*18:01:01, HLA-B*52:01:01 and HLA-C*03:04:01. Four others reached 80% to below 95%, without reaching 95% in an eligible group: HLA-A*30:02:01, HLA-B*35:03:01, HLA-C*07:02:01 and HLA-C*05:01:01. The same 3-field allele could therefore be more readily predictable in one ancestry background yet remain split between competing sequence-groups in another, and global uncertainty often reflected pooled averaging over super-populations with different underlying composition. The adequacy of 3-field class I typing thus depends not only on the allele but on the ancestral composition of the cohort being typed.

### 3.3 Overview of the HLA and MIC allele landscape

Analyses of HLA and MICA/MICB alleles showed that each super-population is characterized by a distinct combination of broadly shared, enriched, and locally restricted alleles. This overall pattern is consistent with previous analyses of the 1000 Genomes panel and broader worldwide HLA surveys, which likewise showed strong ancestry-associated structuring of HLA frequencies rather than a single common global configuration (4, 42, 46). The quantitative locus-level summaries in Figure 3 and Tables 2A and 2B support the interpretation. With HLA labels retained as exported and MIC types summarized at up to two fields, the allele landscape contained 206 common alleles across the 13 analyzed loci. Within this full set, a 137-allele focus set was identified consisting of common alleles that met at least one prioritization criterion: Cramér’s V ≥ 0.20 or population-private or enrichment ratio ≥ 3.0 (focus-set membership is marked in Supplementary Table 2). The largest common-allele catalogs were observed for HLA-B (30), DRB1 (30), HLA-A (24), and HLA-C (24), while DRB3, DRB4, DRB5, and MICB contributed the smallest recurrent allele sets (Table 2A). Together, these summaries indicate that the classical class I loci, particularly HLA-B, contribute much of the visible complexity in the global landscape, whereas the smaller secondary catalogs are concentrated in the DRB paralogues and MICB. The prominence of HLA-B, followed by HLA-A and HLA-C, is concordant with those surveys.

**Figure 3.**
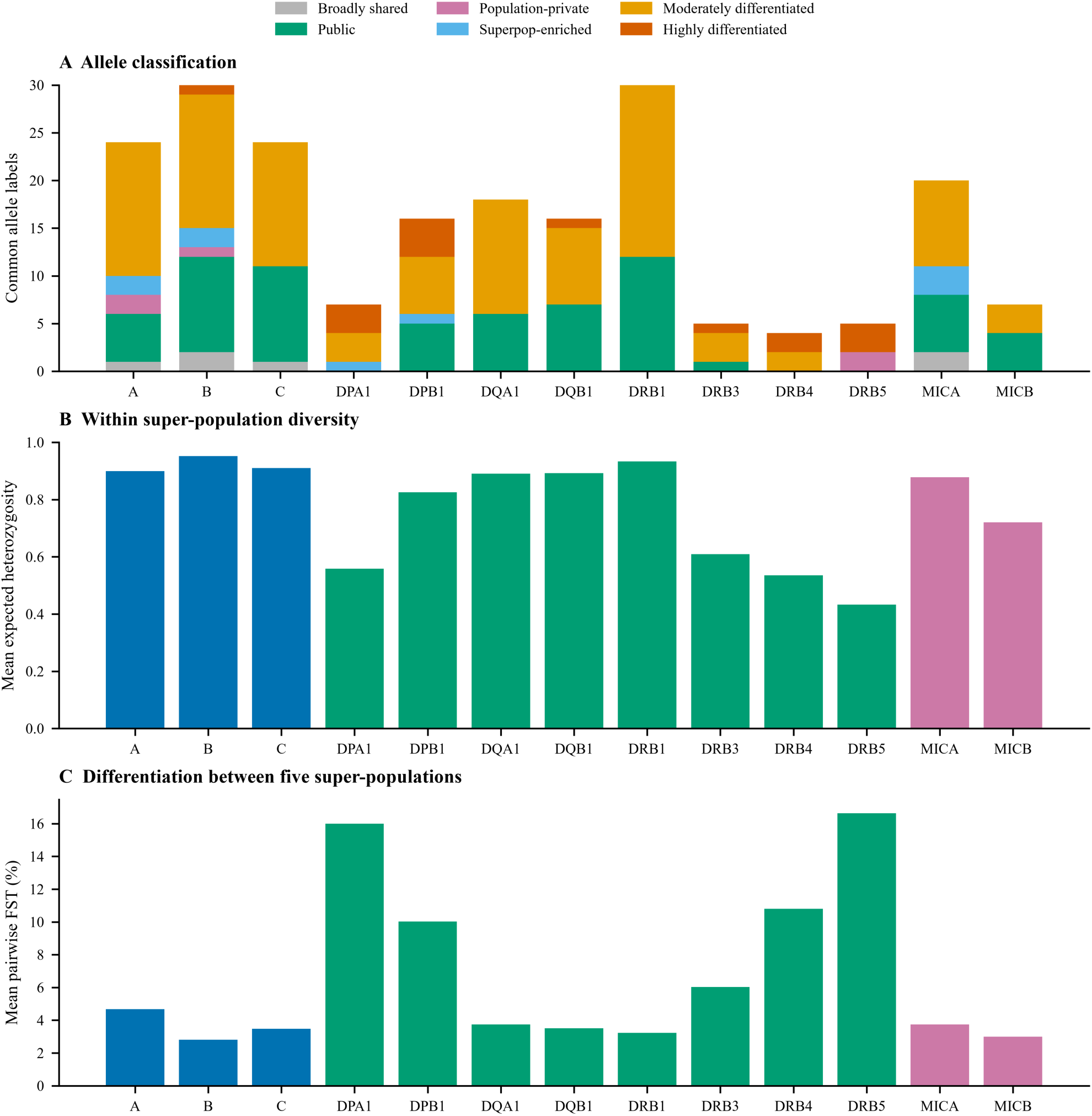
The HLA and MIC allele landscape. (A) Counts of common types (global frequency ≥1%; exported HLA labels and up to two reported MIC fields) by locus and hierarchical classification. Population-private indicates observed presence in one super-population. Public denotes presence in all five groups with Cramér’s V <0.15; it does not require a 1% frequency within every group. (B) Mean expected heterozygosity across five super-populations, calculated from all observed alleles. (C) Mean pairwise Hudson-type FST across the ten super-population pairs, using all observed alleles. DRB3/4/5 metrics are conditional on observed paralogue copies. Top allele identities and frequencies are given in Table 2B.

**Table 2A.** Common allele classification by locus.

| **Locus** | **Common** | **Private** | **Highly**<br>**differentiated** | **Public** | **Moderately**<br>**differentiated** | **Broadly**<br>**shared** | **Group**<br>**enriched** |
| --- | --- | --- | --- | --- | --- | --- | --- |
| A | 24 | 2 | 0 | 5 | 14 | 1 | 2 |
| B | 30 | 1 | 1 | 10 | 14 | 2 | 2 |
| C | 24 | 0 | 0 | 10 | 13 | 1 | 0 |
| DPA1 | 7 | 0 | 3 | 0 | 3 | 0 | 1 |
| DPB1 | 16 | 0 | 4 | 5 | 6 | 0 | 1 |
| DQA1 | 18 | 0 | 0 | 6 | 12 | 0 | 0 |
| DQB1 | 16 | 0 | 1 | 7 | 8 | 0 | 0 |
| DRB1 | 30 | 0 | 0 | 12 | 18 | 0 | 0 |
| DRB3 | 5 | 0 | 1 | 1 | 3 | 0 | 0 |
| DRB4 | 4 | 0 | 2 | 0 | 2 | 0 | 0 |
| DRB5 | 5 | 2 | 3 | 0 | 0 | 0 | 0 |
| MICA | 20 | 0 | 0 | 6 | 9 | 2 | 3 |
| MICB | 7 | 0 | 0 | 4 | 3 | 0 | 0 |

Locus-level diversity metrics further showed that allelic richness and frequency evenness varied substantially across loci. Expected heterozygosity was highest for HLA-B, DRB1, HLA-C, and HLA-A, with mean values of 0.952, 0.934, 0.911, and 0.899, respectively. These values indicate broad and relatively even allele distributions at the major classical loci, consistent with extensive within-population diversity. By contrast, DPA1, DRB4, DRB3, and DRB5 showed lower mean expected heterozygosity, at 0.558, 0.535, 0.609, and 0.433, respectively, consistent with more restricted or more uneven recurrent-allele profiles at these loci (Table 2B). This general contrast is biologically plausible and broadly agrees with earlier work showing that HLA diversity is unevenly distributed across loci and that DRB3, DRB4, and DRB5 differ from the major classical loci because of their DRB1-linked organization and characteristic haplotypic associations (47, 48).

Population differentiation was also strongly locus dependent. The highest locus-level mean FST values were observed for DPA1, DRB5, DRB4, DPB1 and DRB3, with values of 16.0%, 16.6%, 10.8%, 10.0% and 6.0%, respectively (Table 2B). Thus, although HLA-A, HLA-B, HLA-C, and DRB1 contributed the largest common-allele catalogs and the highest within-population diversity, the strongest between-super-population differentiation was concentrated in selected class II and DRB-linked loci.

**Table 2B.** Diversity differentiation and leading alleles.

| **Locus** | **Individuals** | **Mean He** | **Mean FST (%)** | **Leading allele** | **Frequency (%)** |
| --- | --- | --- | --- | --- | --- |
| A | 2,486 | 0.899 | 4.7 | A*02:01:01 | 13.1 |
| B | 2,486 | 0.952 | 2.8 | B*35:01:01 | 6.3 |
| C | 2,486 | 0.911 | 3.5 | C*04:01:01 | 14.3 |
| DPA1 | 2,486 | 0.558 | 16.0 | DPA1*01:03:01 | 48.6 |
| DPB1 | 2,486 | 0.826 | 10.0 | DPB1*04:01:01 | 18.7 |
| DQA1 | 2,486 | 0.890 | 3.7 | DQA1*01:02:01 | 16.9 |
| DQB1 | 2,486 | 0.892 | 3.5 | DQB1*03:01:01 | 13.5 |
| DRB1 | 2,486 | 0.934 | 3.2 | DRB1*07:01:01 | 10.2 |
| DRB3 | 1,569 | 0.609 | 6.0 | DRB3*02:02:01 | 54.2 |
| DRB4 | 1,111 | 0.535 | 10.8 | DRB4*01:03:01 | 60.7 |
| DRB5 | 776 | 0.433 | 16.6 | DRB5*01:01:01 | 66.2 |
| MICA | 2,393 | 0.878 | 3.7 | MICA*002:01 | 19.9 |
| MICB | 2,393 | 0.720 | 3.0 | MICB*005:02 | 43.6 |

### 3.4 Highly differentiated and population-private alleles

Figure 4 and Table 3A summarize allele differentiation across super-populations using Cramér’s V. In this setting, Cramér’s V provides a measure of how unevenly an allele is distributed across groups. Lower values indicate that an allele occurs at relatively similar frequencies across super-populations, whereas higher values indicate progressively stronger concentration in one or a subset of groups. Accordingly, this metric provides a more informative measure of population differentiation than statistical significance alone, because it reflects the magnitude rather than simply the detectability of allele-frequency differences.

**Figure 4.**
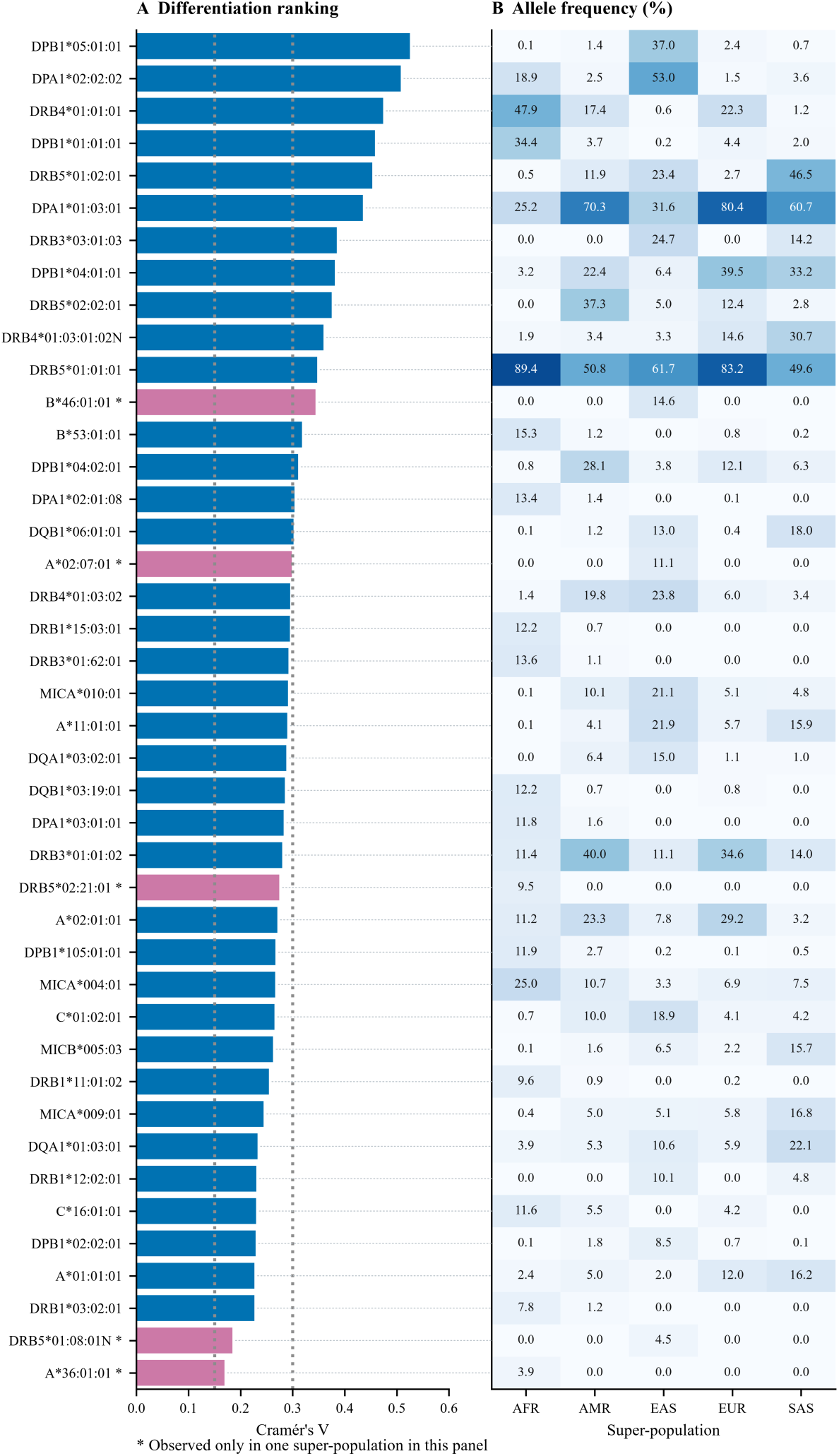
Ancestry informative alleles. (A) Forty common alleles ranked by Cramér’s V, supplemented by population-private alleles outside the ranking. Asterisks and magenta bars mark the five private alleles; vertical dotted lines mark V = 0.15 and 0.30. (B) Frequencies (%) in AFR, AMR, EAS, EUR and SAS for the same alleles. Faint horizontal guides connect each bar to its corresponding heatmap row. Zero denotes no observed copies; denominators are locus-specific. Exact leading values and the private subset are reported in Table 3.

**Table 3A.** Leading differentiated alleles.

| Allele | Cramér's V | Leading group | Pooled frequency (%) | Enrichment ratio |
| --- | --- | --- | --- | --- |
| DPB1*05:01:01 | 0.525 | EAS | 8.8 | 31.4 |
| DPA1*02:02:02 | 0.508 | EAS | 18.2 | 8.0 |
| DRB4*01:01:01 | 0.474 | AFR | 14.9 | 4.6 |
| DPB1*01:01:01 | 0.458 | AFR | 11.4 | 13.4 |
| DRB5*01:02:01 | 0.453 | SAS | 22.2 | 4.8 |
| DPA1*01:03:01 | 0.435 | EUR | 48.6 | 1.7 |
| DRB3*03:01:03 | 0.385 | EAS | 7.3 | 6.9 |
| DPB1*04:01:01 | 0.381 | EUR | 18.7 | 2.4 |
| DRB5*02:02:01 | 0.375 | AMR | 6.3 | 7.4 |
| DRB4*01:03:01:02N | 0.359 | SAS | 11.9 | 5.3 |

The focus-allele analysis identified the specific alleles contributing most strongly to between-super-population differentiation. The strongest-ranked signal was DPB1*05:01:01, which showed a Cramér’s V of 0.525. DPA1*02:02:02 also showed marked continental structure, with a Cramér’s V of 0.508. Additional highly differentiated alleles included DRB4*01:01:01, DPB1*01:01:01, DRB5*01:02:01, DPA1*01:03:01, DRB3*03:01:03, DPB1*04:01:01, DRB5*02:02:01, and DRB4*01:03:01:02N, with Cramér’s V values ranging from 0.474 to 0.359 (Table 3A). Together, these values indicate strong population differentiation, with the highest-ranked alleles showing pronounced frequency concentration in particular super-populations rather than broad frequency balance across all groups.

These results show that the most ancestry-informative alleles were not limited to globally rare or strictly private variants. Several top-ranked alleles remained detectable across multiple super-populations but differed sharply in relative frequency. For example, DPA1*02:02:02 was broadly observed but strongly enriched in EAS, whereas DRB4*01:01:01 and DPB1*01:01:01 were more strongly enriched in AFR. This pattern indicates that super-population structure arises not only from alleles restricted to a single region, but also from major frequency shifts in alleles that remain globally distributed. The enrichment of DPA1*02:02:02 and DPB1*05:01:01 is consistent with their high frequencies reported at three-field resolution in Chinese Han families (49).

Among the classical class I loci, HLA-B*46:01:01 was the clearest population-restricted signal within the differentiated-allele set. This allele was strongly concentrated in EAS and was not observed in the other four super-populations. HLA-A*02:07:01 likewise showed prominent EAS enrichment. These class I examples provide recognizable ancestry-associated markers, but the broader ranked list indicates that class II and DRB-linked variation contributed equally important, and in some cases stronger, population differentiation.

Population-private alleles are summarized in Table 3B. Under the common-allele threshold used for the main landscape analysis, only five population-private alleles were retained: A*36:01:01 and DRB5*02:21:01 in AFR, and A*02:07:01, B*46:01:01 and DRB5*01:08:01N in EAS. DRB5*02:21:01 was observed in 19 copies, representing 2.2% of all observed DRB5 copies and 9.5% of AFR DRB5 copies. No common population-private alleles met the criteria in AMR, EUR or SAS. Examples of rarer population-private types below the common threshold included DPA1*01:03:07 in SAS (13 copies; pooled frequency 0.26%) and DPA1*02:12:01 in AFR (31 copies; 0.62%). In contrast, DRB3*01:62:01 occurred in AFR (105 copies) and AMR (two copies) and was not population-private. Variant and coverage annotations are supplied in Supplementary Table 7. Taken together, Figures 3–4 and Tables 2–3 show that the HLA and MICA/MICB landscape is shaped primarily by frequency differences among shared alleles, with a small common private subset within this panel.

**Table 3B.** Population private common alleles.

| Allele | Observed group | Pooled frequency (%) | Cramér's V |
| --- | --- | --- | --- |
| B*46:01:01 | EAS | 3.2 | 0.344 |
| A*02:07:01 | EAS | 2.4 | 0.299 |
| DRB5*02:21:01 | AFR | 2.2 | 0.275 |
| DRB5*01:08:01N | EAS | 1.1 | 0.184 |
| A*36:01:01 | AFR | 1.1 | 0.169 |

Published population data are broadly consistent with these regional associations while confirming that private here means private to this panel. A*36:01 and the exact A*02:07:01 type have each been reported at low frequency in European American cohorts (50, 51), B*46:01:01:01 and the related DRB5 null type in Thai cohorts (52, 53), the shared two-field DRB5*02:21 type in a Saudi individual (54, 55), and DPA1*02:12 as a single copy in the Maastricht workshop cohort (56). Thai participants were not included in our panel or its SAS group. These occurrences at shared reported fields do not establish identity at deeper fields, and we use population-private for types observed in one sampled super-population.

### 3.5 Multilocus haplotype structure

The leading EM-estimated haplotypes differed among super-populations in each block (Figure 5A– C). In the class I block, the EAS-leading A*02:07:01∼B*46:01:01∼C*01:02:01 combination had an estimated frequency of 8.7%, whereas the leading AFR haplotype, A*30:01:01∼B*42:01:01∼C*17:01:01, was 3.2%, and the leading EUR combination was A*01:01:01∼B*08:01:01∼C*07:01:01 at 5.7%. Corresponding class I backgrounds have been reported at shared fields in Singapore Chinese donors, African American mother–cord pairs and Italian donors, respectively (57–59), placing differentiated individual alleles within recurrent multilocus backgrounds.

**Figure 5.**
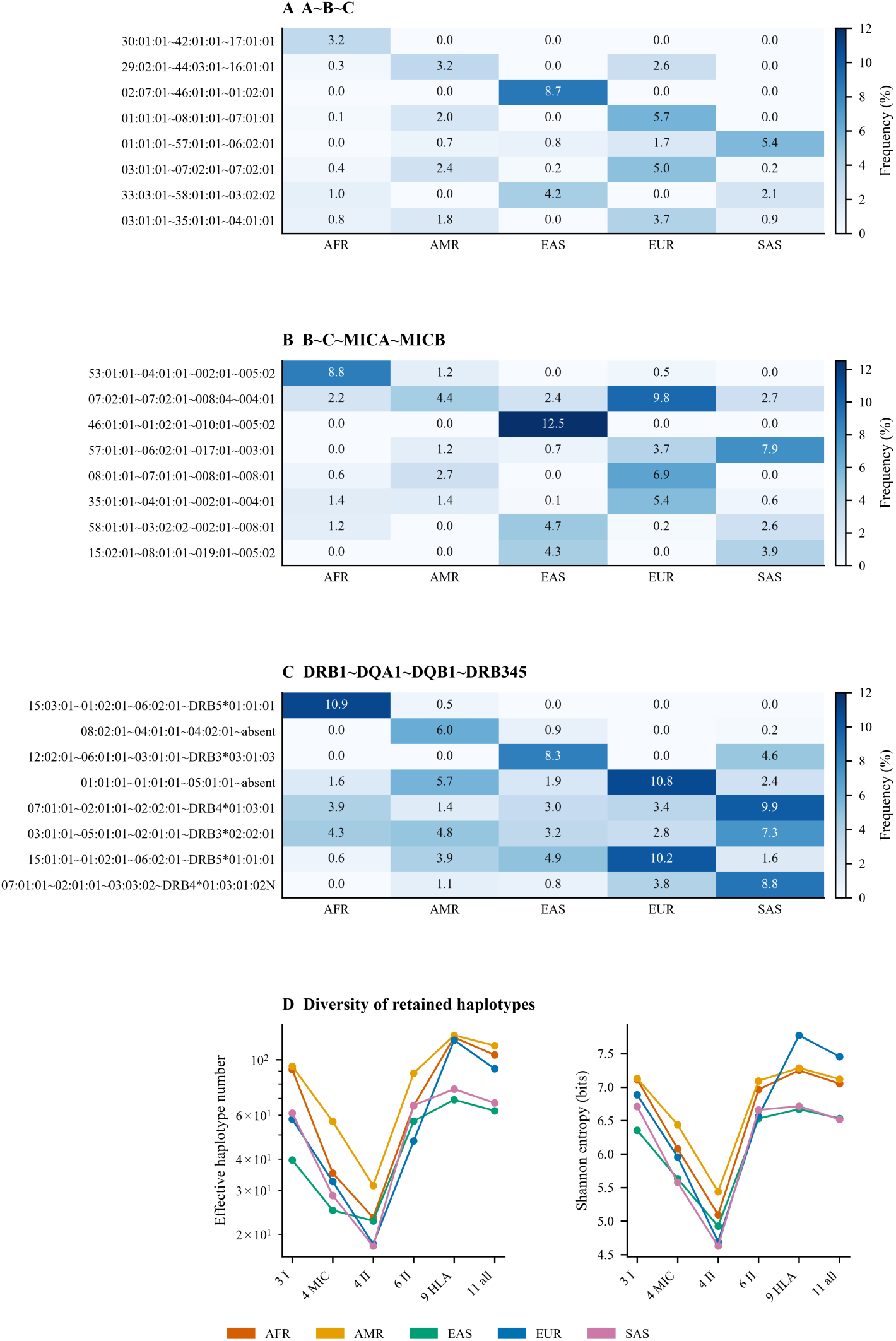
Multilocus haplotype structure. (A–C) EM-estimated frequencies (%) of eight selected haplotypes in the class I, MIC-linked and class II core blocks. Selection retains the leading haplotype in each super-population and fills remaining rows by unweighted mean frequency. Alleles follow the locus order in each panel title; identical retained MIC haplotypes were combined at two reported fields. Complete identities are given in the haplotype keys. DRB345 represents the exported DRB3/4/5 call or an absence state. (D) Effective number (log scale) and Shannon entropy in bits calculated after normalizing all retained haplotype frequencies within each block and group. Labels indicate locus count and block. These are conditional diversity summaries; retained mass and sample sizes are reported in Supplementary Table 3. Connecting lines guide comparison and do not represent a common nested sequence of blocks.

The MIC-linked block preserved a distinct extension of the class I structure. B*46:01:01∼C*01:02:01∼MICA*010:01∼MICB*005:02 was the leading EAS combination at 12.5%, while B*53:01:01∼C*04:01:01∼MICA*002:01∼MICB*005:02 led AFR at 8.8%. B*07:02:01∼C*07:02:01∼MICA*008:04∼MICB*004:01 led both AMR and EUR, at 4.4% and 9.8%, respectively. The B∼C∼MICA∼MICB block is the bridge between the population resource and MIC prediction: its estimated associations provide the frequency model used to recover MIC types from HLA-B/C genotypes. Prediction accuracy was assessed separately with training-only frequencies in each cross-validation fold (Figure 6), rather than inferred from the retained full-panel haplotype heatmap.

**Figure 6.**
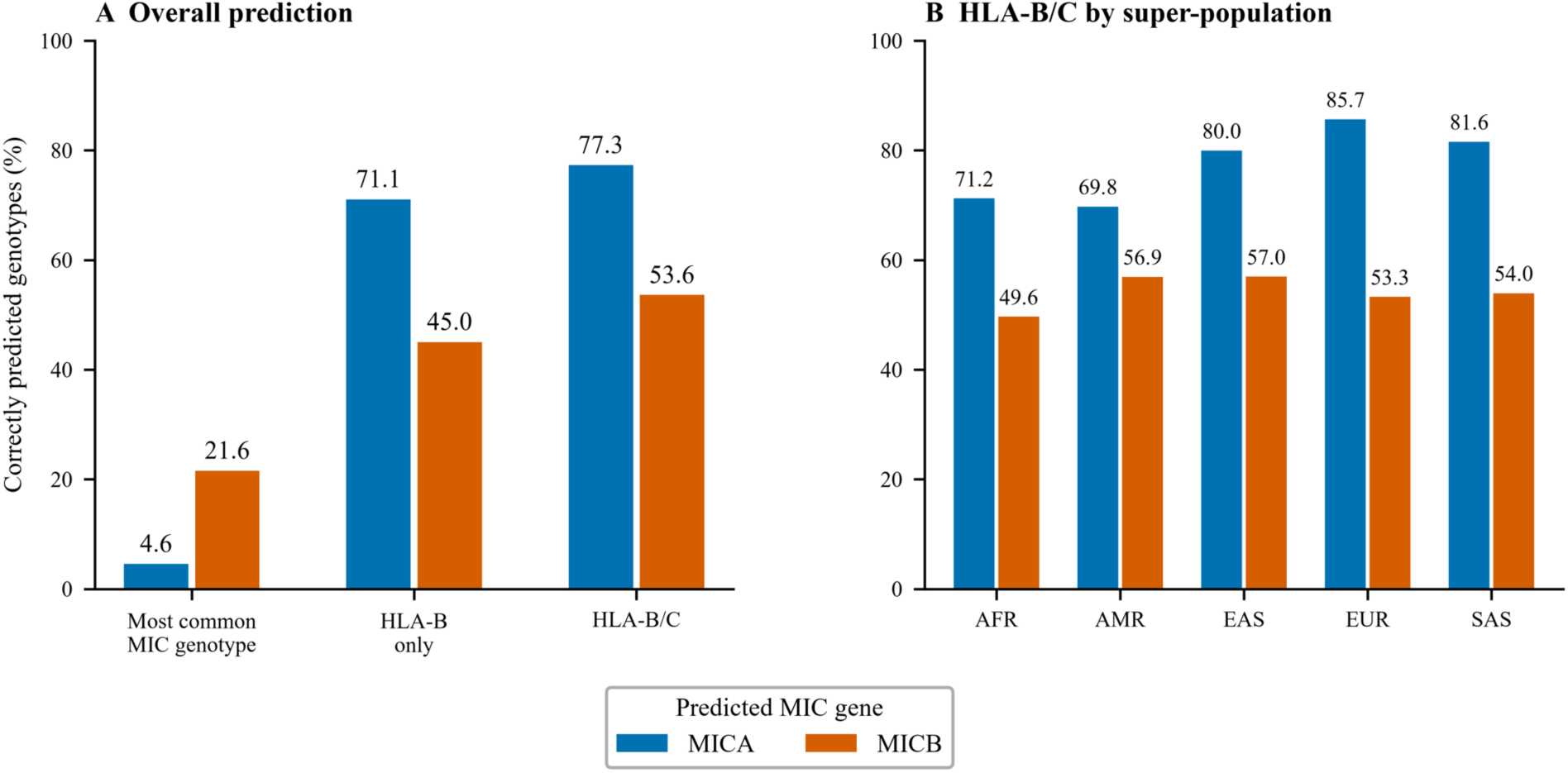
Prediction of MICA and MICB from HLA types. (A) Exact genotype accuracy using the most frequent training MIC genotype without HLA input (majority baseline), two-field HLA-B alone, and two-field HLA-B/C. (B) Two-field HLA-B/C accuracy by super-population, using pooled training frequencies. A correct genotype requires both MIC alleles to match, irrespective of order. Blue bars denote MICA and orange bars MICB; numbers show percentages. Each configuration was evaluated in the same 2,393 individuals by five-fold stratified cross-validation, with haplotype frequencies estimated from training individuals only. MIC targets use up to two reported numeric fields, retaining expression suffixes and shorter or provisional labels; third-field intronic extensions are excluded. All unsupported-input and rare-allele predictions remain in the denominator. Intervals, other configurations, group baselines, confidence analyses and reference diagnostics are supplied in Supplementary Table 4.

Class II core haplotypes were comparatively concentrated among the retained estimates. Conditional effective haplotype numbers ranged from 17.9 to 31.3, compared with 39.6–94.2 for class I and 24.9–56.6 for the MIC-linked block (Figure 5D; Supplementary Table 3). The retained frequency mass was 91.4–94.6% for class II core, 73.0–89.8% for class I and 81.9–96.0% for the MIC-linked block. The main figure retains class I, MIC-linked and class II core haplotypes. The detailed extended class II and eleven-locus combinations are supplementary because their high diversity and incomplete retained frequency mass limit interpretation (Supplementary Figures 2 and 3). In the eleven-locus block, the leading estimates ranged from 1.2% to 3.2%, but only 26.1–64.2% of frequency mass was retained. Consequently, extended-block diversity cannot be inferred from the exported summaries without accounting for the omitted haplotypes.

### 3.6 Prediction of MICA and MICB from HLA type

Two-field HLA-B/C input predicted MICA genotypes correctly in 1,850 of 2,393 held-out individuals, an accuracy of 77.3% (95% interval, 75.6–78.9%), compared with 4.6% for the training-derived majority-genotype baseline. MICB accuracy was 53.6% (1,283 of 2,393; 95% interval, 51.6–55.6%), compared with a 21.6% baseline (Figure 6A; Supplementary Table 4). Using HLA-B alone gave accuracies of 71.0% for MICA and 45.0% for MICB. Adding HLA-C therefore improved prediction for both genes, while MICA remained more predictable than MICB. HLA-B/C accuracy varied across super-populations, ranging from 69.8% to 85.7% for MICA and 49.6% to 57.0% for MICB (Figure 6B). Population-specific frequencies and three-field HLA-B/C input gave smaller overall gains than adding HLA-C (Supplementary Table 4). Allele-level performance and confidence analyses are provided in Supplementary Figures 4 and 5. These internally cross-validated results demonstrate recoverable MIC information from HLA types, with incomplete and population-dependent accuracy.

## 4 Discussion

This study extends the 1000 Genomes Phase 3 panel to high-resolution HLA and MICA/MICB typing and uses that resource to address the three questions set out above. The sections below consider the quality of the resource, what extended class I resolution does and does not add, the locus-specific structure of the allele and haplotype landscape, and how far MIC variation can be recovered from HLA type.

### 4.1 Resource quality and concordance with previous typing

The quality of the typing resource was assessed in three ways: control and duplicate samples distributed across plates, comparison with previous typing of shared 1000 Genomes individuals, and independent retyping of discrepant samples. Direct comparison is more limited for DRB3, DRB4, DRB5, MICA and MICB than for HLA-A, HLA-B, HLA-C or DRB1, because earlier datasets often focused on the classical transplantation loci and did not analyze the other genes at comparable scale and resolution. The two-field comparison distinguishes explicit agreement from compatibility with listed ambiguous genotypes, and neither dataset is treated as a reference truth set.

The twenty reciprocal five-locus cross-individual profile matches between the present study and the 2018 release raised the question of whether genotypes and sample identifiers corresponded correctly. To investigate these discrepancies, the affected samples (n = 20) were independently re-obtained from the original repository and retyped. The repeat genotyping reproduced the sample assignments reported here, providing independent support for their accuracy and indicating that the reciprocal discrepancies originated in the earlier dataset, although their precise cause remains undetermined (Table 4; Supplementary Tables 5 and 6).

**Table 4.** Direct comparison of recorded HLA types. Reported genotypes are compared at two-field resolution, retaining expression suffixes. Explicit matches and matches compatible with listed alternatives are counted separately. The two allele copies are compared without regard to order.

| Locus | Individuals compared | Explicit matches | Ambiguity matches | Different types | Match (%) |
| --- | --- | --- | --- | --- | --- |
| HLA-A | 962 | 131 | 783 | 48 | 95.0 |
| HLA-B | 962 | 194 | 724 | 44 | 95.4 |
| HLA-C | 961 | 103 | 797 | 61 | 93.7 |
| HLA-DRB1 | 962 | 401 | 496 | 65 | 93.2 |
| HLA-DQB1 | 960 | 362 | 519 | 79 | 91.8 |

**B Current study versus Abi-Rached et al. (43)** 2,409 individuals have identifiers present in both datasets.
| Locus | Individuals compared | Explicit matches | Ambiguity matches | Different types | Match (%) |
| --- | --- | --- | --- | --- | --- |
| HLA-A | 2,409 | 2,307 | 31 | 71 | 97.1 |
| HLA-B | 2,403 | 2,302 | 15 | 86 | 96.4 |
| HLA-C | 2,407 | 2,331 | 9 | 67 | 97.2 |
| HLA-DRB1 | 2,408 | 2,343 | 3 | 62 | 97.4 |
| HLA-DQB1 | 2,119 | 2,010 | 28 | 81 | 96.2 |

Explicit matches: both studies report one matching allele pair. Ambiguity matches: at least one study lists alternatives, and one complete pair of listed alleles matches the other genotype. Both copies must match, retaining homozygous multiplicity.

### 4.2 Fourth-field variation: sequence boundaries, biological significance, and practical utility

The interpretation of HLA fourth-field diversity requires consideration of the genomic sequence boundaries used to define individual alleles. Whereas three-field HLA designations distinguish alleles according to their encoded protein sequence and synonymous coding variation, fourth-field designations identify differences outside the coding sequence. Consequently, the number of distinguishable fourth-field variants within a three-field lineage depends partly on the extent of noncoding sequence available for comparison. As additional intronic and flanking sequences are characterized, further variants may be distinguished without any corresponding change in the encoded HLA protein. The catalogue-to-cohort comparison reported above illustrates why catalogued fourth-field counts should not be equated with biologically meaningful diversity.

Our analysis addressed the dependence on sequence boundaries by defining sequence groups within the genomic intervals covered by the typing assays, rather than treating every catalogued fourth-field designation as independently distinguishable. Within these standardized boundaries, most common three-field class I lineages were dominated by a single extended sequence group. Across the cohort, 91.8% of assigned allele copies fell within their predominant within-lineage group, and the 8.2% that did not were concentrated in a small number of recurrent binary splits rather than distributed across the catalogued fourth-field range. At least one such distinction was present in 39.5% of individuals, but that proportion follows from the per-copy rate rather than from a correspondingly broad reservoir of variation: with two copies at each of three loci, independent sampling at the observed per-copy rates would itself yield approximately 40%. The variation that extended typing resolves in this panel is therefore limited and repetitive, and its frequency of occurrence should not be confused with its biological or clinical significance.

Fourth-field assignments are important when their underlying differences influence HLA expression, RNA processing, or other functional properties. The genomic regions interrogated by the standard ScisGo-HLA-v6 assay encompass all currently recognized fourth-field variants demonstrated to have such consequences, as documented in the kit supporting materials. Thus, the assay retains established functionally relevant variation without requiring resolution of every additional noncoding distinction represented in more extensive genomic reference sequences. In transplantation, a fourth-field mismatch does not itself establish an effect on allorecognition, graft-versus-host disease, or other clinical outcomes. Similarly, many rare fourth-field variants occur too infrequently to support adequately powered population association analyses individually. Increasing the amount of noncoding sequence examined may therefore expand the number of named alleles without a corresponding increase in established clinical or research utility.

Importantly, eleven common three-field lineages exhibited substantial heterogeneity within the defined typing intervals, with their predominant sequence groups representing less than 80% of assigned copies. The distributions of the leading groups differed among ancestral populations, and the distinguishing sequence differences involved intronic regions, the 3′ untranslated region, or both. Although the functional consequences of these particular distinctions remain unknown, their recurrent occurrence and population-specific distributions make them more tractable candidates for experimental investigation than an unrestricted collection of rare fourth-field alleles.

Together, these findings indicate that the value of extended HLA typing should be assessed according to the biological information it provides rather than the number of additional allele designations it produces. Standardized typing boundaries establish a reproducible measure of sequence diversity, while targeted functional investigation of recurrent noncoding variants offers a focused opportunity to identify additional HLA variation of biological or clinical significance.

### 4.3 Locus-specific diversity and population differentiation

One of the clearest contrasts in these data was between the loci carrying the most allelic diversity and those carrying the most ancestry information. HLA-B, DRB1, HLA-A and HLA-C held the largest common-allele catalogs and the highest expected heterozygosity (0.899–0.952), whereas the highest mean FST values fell on DRB5 (16.6%), DPA1 (16.0%), DRB4 (10.8%), DPB1 (10.0%) and DRB3 (6.0%).

Part of this contrast follows from the metric rather than the biology. Differentiation estimates depend on allele-frequency distributions and within-population diversity, so comparisons across loci should be interpreted with the estimator and resolution held in view. High diversity and comparatively modest FST at the classical loci should not be read as evidence that their allele frequencies are geographically uniform (60).

The loci showing elevated differentiation also share features that make large frequency differences expected. DRB3, DRB4 and DRB5 are not present on every haplotype: they generally associate with the DR52, DR53 and DR51 haplotype groups, respectively, although unusual associations and null alleles occur (61). Here, their allele frequencies and differentiation were calculated conditional on observed paralogue copies. These estimates describe variation among carriers and do not directly measure the frequencies of paralogue absence or DR haplotype groups in the entire cohort. DRB5 differentiation must therefore be interpreted within that conditional denominator. HLA-DPA1 and HLA-DPB1 differ for a separate reason: recombination hotspots between the DR-DQ and DP regions reduce their linkage, although this does not establish complete independence in every population (62). Both sets of loci also have small common-allele catalogs dominated by a single allele — four to seven common alleles at DPA1, DRB3, DRB4 and DRB5, with the most frequent accounting for 49% to 66% of chromosomes — and frequency shifts in these dominant alleles can strongly affect diversity and differentiation summaries.

The practical consequence is that the most ancestry-informative markers in this panel were not the loci that dominate the visible complexity of the landscape. The focus-allele analysis made the same point at allele level, with the strongest signals at DPB1*05:01:01 and DPA1*02:02:02 rather than at any class I allele. This bears on the transplant setting, where HLA-DPB1 mismatching is already recognized as a determinant of outcome and DRB3/4/5 matching is increasingly considered: between-population frequency shifts can alter the availability of matching alleles, although differentiation values do not by themselves rank absolute donor–recipient mismatch probabilities. This pattern of strong geography-associated structure alongside marked locus-to-locus heterogeneity is concordant with earlier HLA population-genetic studies (42, 47).

### 4.4 Multilocus HLA and MIC haplotypes

Multilocus haplotype analyses extended the population-specific patterns observed at individual HLA and MIC loci, revealing characteristic allele combinations that differed in frequency and diversity among ancestral populations. The HLA-A*02:07:01∼B*46:01:01∼C*01:02:01 haplotype was the most frequent class I combination in EAS (8.7%), whereas A*30:01:01∼B*42:01:01∼C*17:01:01 predominated in AFR (3.2%) and A*01:01:01∼B*08:01:01∼C*07:01:01 in EUR (5.7%). Extending these analyses to MICA and MICB identified similarly structured HLA-B∼C∼MICA∼MICB haplotypes, including B*46:01:01∼C*01:02:01∼MICA*010:01∼MICB*005:02 in EAS (12.5%) and B*53:01:01∼C*04:01:01∼MICA*002:01∼MICB*005:02 in AFR (8.8%). These associations demonstrate that MIC alleles occur within recognizable HLA haplotypic backgrounds, providing the population-genetic rationale for predicting MICA and MICB genotypes from HLA-B and HLA-C typing. The predictive performance of these associations was evaluated separately using cross-validation, with haplotype frequencies estimated from training samples.

The greater concentration of retained class II core haplotypes relative to class I combinations further illustrates differences in multilocus diversity across the MHC. Haplotype frequencies were estimated statistically from unphased genotypes and rare combinations were excluded before summary, so the estimates are conditional on the retained distribution; within that limit they establish a reference for investigating HLA–MIC associations across populations.

### 4.5 MICA/MICB variation and prediction from HLA genotypes

MICA and MICB encode ligands of the activating NKG2D receptor, linking their variation to immune recognition of stressed and transformed cells. The MICA A5.1 group, whose prototype is MICA*008, contains a transmembrane-region insertion associated with a truncated, GPI-anchored protein, illustrating why MIC allele variation can have biological consequences (63). Associations of NKG2D ligand and receptor variation with transplantation outcomes further motivate joint characterization of these loci (64).

Prediction uses the nonrandom association of HLA and MIC alleles on shared haplotypes. Such associations have been described for HLA-B, MICA and MICB, including HLA-B–MICA haplotypes associated with more than one MICB allele (31). Cross-validation measures how often these associations recover a complete MIC genotype in held-out individuals; it is not a measure of linkage disequilibrium strength, so the greater MICA accuracy does not by itself establish stronger LD than for MICB.

The exact-genotype accuracies achieved with two-field HLA-B/C typing (77.3% for MICA and 53.6% for MICB) indicate that existing HLA records can recover substantial MIC information, but the distinction between research utility and reliable individual-level inference is important. Prediction errors may vary with population background, MIC allele frequency, or haplotypes insufficiently represented in the reference. Retrospective association studies using predicted MIC genotypes would therefore need to account for uncertainty and evaluate possible attenuation or bias of estimated effects, and direct MIC typing remains preferable where accurate individual-level assignment is essential. Validation in independent cohorts, and analyses of prediction confidence and accuracy by allele frequency, would help define appropriate applications.

### 4.6 Practical implications of the integrated typing approach

The typing approach has practical implications for assembling datasets of this kind. The HLA and MICA/MICB assays follow the same two-stage amplicon workflow and differ only in their primer content, so the two gene systems could in principle be combined into a single reaction set yielding high-resolution typing for both from one application. Library preparation consisted of DNA and buffer addition, pooling, purification and loading, with no sample transfer between wells, and the complete 13-locus panel was generated on a benchtop sequencing platform. Cost and throughput were not benchmarked against long-range PCR shotgun approaches.

### 4.7 Applications to MHC association studies and transplantation cohorts

This dataset is a resource for the many published GWAS implicating the MHC. The region contains numerous disease-associated signals, but identifying the causal genes and alleles remains difficult where haplotypic association is extensive (65). Combined with the existing high-coverage whole-genome sequencing data available for the 1000 Genomes samples, the HLA and MICA/MICB allele calls generated here provide a foundation for constructing higher-resolution imputation reference panels for the MHC. Such panels could support fine-mapping of existing disease associations using individual-level genotype data or appropriate summary-statistics-based methods, potentially helping to distinguish associations attributable to classical HLA alleles, MIC variation, or other linked MHC variants. The MIC predictor reported above addresses a complementary and more immediately accessible application: because it takes HLA type rather than SNP genotypes as input, it can be evaluated in transplant cohorts and donor registries that already hold HLA typing without MIC typing, and the internal cross-validation quantifies how much MIC variation is recoverable within this reference. Independent-cohort validation remains necessary.

### 4.8 Limitations of the haplotype and prediction analyses

The haplotype estimates are inferred from unphased genotypes and are sensitive to sample size, rare-allele pooling and the number of loci considered. In particular, the analytical export removed low-frequency and pooled-rare haplotypes before the diversity summaries were generated. Normalizing the retained frequencies corrects the mathematical calculation of conditional diversity, but it cannot recover complete-block diversity or resolve potential sensitivity to the copy-specific pooling procedure.

The MIC evaluation used random folds within the paired reference panel and did not test a separately collected cohort. Relatedness was not assessed, and the folds distributed individuals rather than pedigrees. The narrow retained MIC sample from IBS and uneven group sizes further limit population comparisons. Training-unseen alleles, unsupported HLA combinations and all fallback calls remained in overall accuracy. The released full-panel haplotypes support reproducibility but do not independently validate the predictor outside this dataset. Original third-field MIC intronic extensions and provisional tags remain traceable in the released source-annotation data. Sequence-localization annotations in Table 1C retain their reported reference-position convention and require the original pairwise sequence alignments for independent reconstruction. HLA variant and coverage annotations are retained verbatim in Supplementary Table 7.

### 4.9 Conclusions

This study provides a high-resolution HLA and MICA/MICB reference resource for the 1000 Genomes Phase 3 panel and demonstrates the importance of distinguishing extended sequence resolution from biologically meaningful HLA variation. Most common three-field class I lineages were dominated by a single sequence group within defined typing boundaries, whereas eleven lineages exhibited substantial heterogeneity involving recurrent noncoding differences. Although additional sequence discrimination was observed in a substantial fraction of individuals, its clinical relevance remains unestablished. The standard typing assay includes currently recognized functionally consequential fourth-field variants, while the recurrent noncoding differences identified through extended typing provide focused candidates for future functional investigation.

The resource also reveals population-associated differences in HLA and MIC allele frequencies and haplotypes and demonstrates that existing HLA-B/C genotypes can recover MICA and MICB genotypes with cross-validated accuracies of 77.3% and 53.6%, respectively. These findings provide a foundation for investigating MHC-associated disease variation and for evaluating the contribution of MIC polymorphism to transplantation outcomes. Further functional studies and independent validation will be necessary to establish the clinical utility of the additional sequence distinctions and predicted MIC genotypes.

## Supporting information

all supplementary files

## Data availability statement

The allele-call data and code supporting the analyses in this study are provided in the accompanying data and code package. The package contains original typing-call exports and previously published comparison datasets, processed HLA and MIC genotypes with population annotations, allele-frequency and haplotype estimates, MIC prediction results and reference models, and the scripts used for analysis and figure and table generation. Supplementary Tables 1–7 and Supplementary Figures 1–6 provide the principal supporting results. A README, data dictionary and checksums describe the files and support reproducibility. Review codes replace individual identifiers in manuscript displays and supplementary tables; public 1000 Genomes identifiers are retained in the analytical data. Original sequencing reads are not included in this package.

## Ethics statement

This study analyzed publicly available, de-identified DNA samples and genomic data from the 1000 Genomes Project, distributed by the NHGRI Sample Repository for Human Genetic Research at the Coriell Institute for Medical Research. Informed consent for broad data sharing and secondary research use was obtained from participants by the original project. Ethical approval was not required for this secondary analysis in accordance with local legislation and institutional requirements.

## Author contributions

CWP and DEG conceived the study and designed the analyses. CWP and RW developed and performed the HLA and MICA/MICB typing assays, including the supplementary class I intron-targeting reactions. YS, AT, DK, SP, and CWP performed sample handling, plate design and laboratory quality control. WCN, EBG and RW developed the sequence-analysis and allele-assignment pipeline and curated the allele calls. CWP performed the allele-frequency, diversity and differentiation analyses. CWP performed the haplotype estimation and the MICA/MICB prediction analyses. CWP, EBG, AI and RW performed the comparison with earlier 1000 Genomes typing. CWP and DEG prepared the figures and tables. DEG wrote the original draft. All authors reviewed and edited the manuscript and approved the submitted version. DEG supervised the study and acquired funding.

## Acknowledgments

We thank the individuals who donated samples to the 1000 Genomes Project; this work would not have been possible without their participation. We acknowledge the 1000 Genomes Project Consortium and the International Genome Sample Resource (IGSR) at EMBL-EBI for curating and providing continued access to the panel and its associated population data (25, 26), and the NHGRI Sample Repository for Human Genetic Research at the Coriell Institute for Medical Research for distribution of the DNA samples.

## Generative AI statement

During preparation of this manuscript the authors used Claude (Anthropic) and Codex (OpenAI) to assist with drafting and editing of manuscript text, including figure and table legends, to check internal consistency across sections, and to suggest structural and analytical revisions. All laboratory work, data generation, statistical analysis and interpretation of results were performed by the authors. The authors reviewed and verified all content and take full responsibility for the published work.

## Supplementary material

Supplementary Tables 1–7 and Supplementary Figures 1–6 accompany this article and are described in the data availability statement.

