## Supplementary material for "High-resolution HLA and MICA/MICB typing of the 1000 Genomes panel: a resource for precision medicine": all supplementary files: SupplementaryFigure1_Typing_workflow.pdf

### Universal targeted amplicon workflow for HLA and MIC typing

Multiplex PCR → paired-end sequencing → centralized cloud analysis → interactive review → high-resolution allele calls

**1 DNA input**

Genomic DNA enters a coordinated four-reaction assay framework.

**2 Stage 1 PCR**

Four coordinated reactions enrich exon-containing regions across HLA and MIC targets in parallel.

**3 Stage 2 PCR**

Sequencing-compatible products are generated directly from stage 1 amplicons without sample transfer between wells.

**4 Pool / purify / normalize**

Reaction sets are combined, column-purified, quantified, normalized, and prepared for sequencing.

**5 Illumina sequencing**

Libraries are loaded on MiSeq or an equivalent paired-end platform for batch-oriented processing.

**6 Cloud analysis**

Standardized preprocessing, QC, reference matching, allele assignment, traceable reporting, and controlled reanalysis

**7 Review and export**

Desktop or web review supports evidence inspection, targeted editing, final calls, and report-ready export.

**Allele calls**

#### Coverage and phasing features

- Exon-focused sequence evidence is generated across targeted HLA and MIC loci.
- Controls and duplicate samples support laboratory quality control.
- HLA-A, -B and -C retain phasing across exons 1–3, 4–5 and 6–8.
- HLA-DPB1 retains phasing across exons 4–5.
- Standard HLA-A, -B and -C coverage omits intron 3 and includes partial 5′ and 3′ UTR sequence.
- Extended class I typing adds two supplementary intron-targeting reactions.

#### Laboratory and analytical implementation

- Stage 1: 2 µL DNA plus 8 µL S1 buffer.
- Stage 2: add 10 µL S2 buffer to the same wells.
- ScisGo-HLA-v6 and ScisGo-MICAB-v3 assays.
- Paired-end Illumina sequencing of pooled products.
- Cloud analysis and GeMS-UI review support traceable allele reports.
- IPD-IMGT/HLA 3.66.0 (October 2026); typing in 2018–2019.

Extended assignments describe sequence within assay coverage. Candidates distinguished only outside covered regions remain unresolved.
