## Supplementary material for "High-resolution HLA and MICA/MICB typing of the 1000 Genomes panel: a resource for precision medicine": all supplementary files: SupplementaryFigure6_Weighted_resolution.pdf

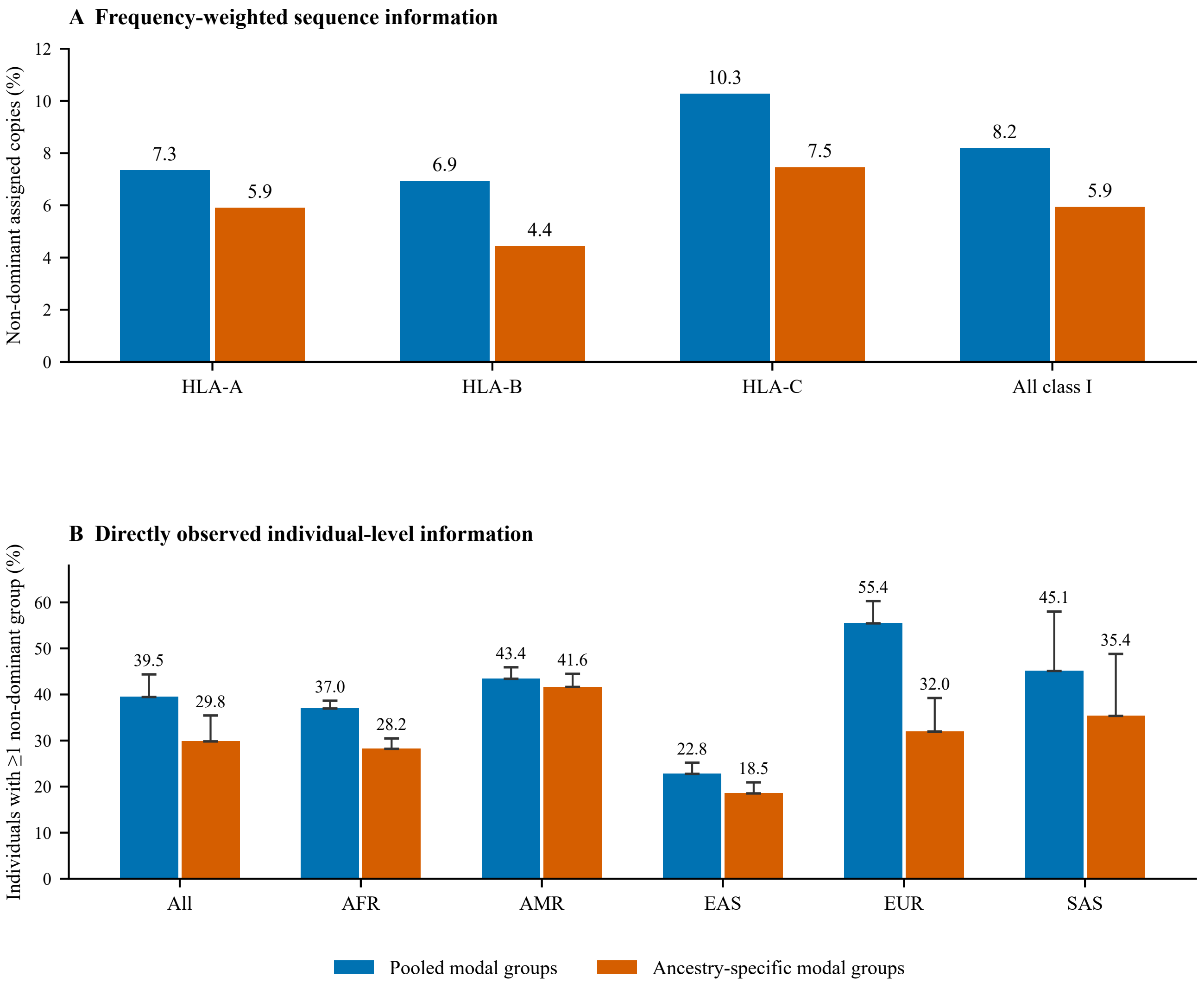


Bars in B: observed individuals. Whiskers: additional unresolved-call bounds, not confidence intervals.
Modal groups estimated in this cohort; descriptive discrimination does not establish clinical benefit.
