## Supplementary material for "High-resolution HLA and MICA/MICB typing of the 1000 Genomes panel: a resource for precision medicine": all supplementary files: SupplementaryFigure_legends.docx

Supplementary figure legends

#### Supplementary Figure 1 Typing workflow

The original seven-stage schematic shows the two-stage multiplex PCR workflow, pooling and purification, paired-end Illumina sequencing, centralized analysis, and review/export. The standard assay uses four reactions; extended HLA-A, -B and -C typing adds two supplementary reactions for intron 3 and partial terminal-region evidence. Typing was performed in 2018–2019 using IPD-IMGT/HLA release 3.66.0 (October 2026). Candidates distinguished only outside assay coverage remain unresolved. The original diagram layout is retained; wording describes assay design and implementation without claiming measured advantages in cost, throughput or allelic-dropout performance.

#### Supplementary Figure 2 Extended class II haplotypes

EM-estimated frequencies (%) for 15 selected haplotypes of DPA1~DPB1~DQA1~DQB1~DRB1~DRB345. The selection rule is the same as Figure 5. Rows use identifiers ExtII-01 to ExtII-15; complete allele strings are in the accompanying haplotype key and Supplementary Table 3 source data. Frequencies are retained EM estimates and do not sum to one across the displayed subset.

#### Supplementary Figure 3 Combined HLA and MIC haplotypes

EM-estimated frequencies (%) for 15 selected haplotypes of the eleven-locus block A~B~C~DPA1~DPB1~DQA1~DQB1~DRB1~DRB345~MICA~MICB. Rows use identifiers TotalHLA-01 to TotalHLA-15 with complete allele strings supplied in the haplotype key. The figure describes leading retained combinations; exported frequency mass and conditional diversity are given in Supplementary Table 3. MIC types use up to two reported fields; identical retained haplotypes were combined at that resolution without changing retained frequency mass.

#### Supplementary Figure 4 MIC per allele performance

Allele-copy sensitivity and positive predictive value for (A) MICA and (B) MICB under primary pooled two-field HLA-B/C prediction of MIC types at up to two fields. Each allele is evaluated using only held-out allele-fold observations for which its training count was at least ten. Right-hand counts show observed copies across eligible folds. The reporting filter does not remove rare-allele carriers or errors from overall genotype accuracy. Primary per-allele summaries are supplied in Supplementary Table 4; complete fold-specific training counts, reporting eligibility and rare or unseen inventories remain in the released MIC evaluation data.

#### Supplementary Figure 5 MIC prediction confidence

(A) Accuracy among accepted calls (solid) and fraction of all individuals receiving a call (dashed) across minimum posterior thresholds for pooled two-field HLA-B/C prediction. (B) Mean model posterior probability versus observed exact-genotype accuracy in fixed bins containing at least ten calls; the dotted diagonal indicates equality. Blue denotes MICA and orange MICB. Confidence filtering reduces the number of reported calls, and model probabilities are not established as calibrated confidence estimates. Primary threshold and calibration-bin summaries are provided in Supplementary Table 4; full group and additional-configuration records remain in the released MIC evaluation data.

### Supplementary Figure 6 Cohort weighted fourth field information

(A) Non-dominant assigned-copy fractions at HLA-A, HLA-B and HLA-C and across all three loci, using pooled or ancestry-specific modal sequence groups. (B) Directly observed proportions of individuals with at least one known non-dominant group across class I. Bars use all individuals in each population; whiskers extend to the upper bound allowing individuals without a known event but with unresolved group assignment to have an event. Whiskers are unresolved-call bounds, not confidence intervals. All reported lineages are included; shorter and four-field primary labels are retained without field completion. Modal groups are estimated in this cohort. Ties use the numerically lowest representative; exhaustive alternative tie counts, common-lineage and explicitly three-field-only sensitivities, and all denominators are supplied in Supplementary Table 1. These are descriptive measures of additional discrimination, not validated prospective predictions or evidence of clinical benefit.
