## Supplementary material for "High-resolution HLA and MICA/MICB typing of the 1000 Genomes panel: a resource for precision medicine": all supplementary files: SupplementaryMethods.docx

Supplementary Methods

### S1 Extended class I typing and sequence groups

The analytical HLA cohort contained 2,486 individuals with extended HLA-A, -B and -C assignments. The typing intervals included the gene bodies and proximal genomic flanks. Relative to the ATG start and stop codons, HLA-A extended 300 bp upstream and 318 bp downstream, HLA-B 284 bp upstream and 586 bp downstream, and HLA-C 319 bp upstream and 394 bp downstream. Canonical comparison intervals were genomic positions 1–3503 of HLA-A*01:01:01:01, 1–3546 of HLA-B*07:02:01:01 and 177–3784 of HLA-C*01:02:01:01. The canonical HLA-A genomic reference contained 300 downstream bases; 18 additional assay bases were not represented in that record. The final supplied assay intervals were used without additional trimming.

Reference sequences were reconstructed from the release-pinned IPD-IMGT/HLA 3.66.0 genomic alignments. A hyphen denoted identity with the alignment reference, a dot an aligned insertion/deletion position, and an asterisk unavailable sequence. Complete sequences with identical gap-free sequence within the interval were grouped. Shorter references were retained at their available positions; missing reference bases were not filled and did not imply missing sample sequencing. Within a three-field lineage, a shorter reference joined a complete group only when exactly one group was compatible at jointly available positions. Otherwise mutually compatible shorter profiles were retained separately. Nontransitive chains of compatibility were not merged. Group display labels favored the numerically lowest complete member, or the lowest shorter member if no complete member was available; this was not an inference of candidate probability (Matern et al., 2021).

Laboratory-confirmed within-assay assignments took precedence over expanded candidate lists where such confirmation was available. For HLA-B*40:02:01, the author confirmed that the :01-versus-:03 difference 264 bp upstream of ATG was callable and that the original sequencing-supported groups were 72 copies represented by :01 and one by :07. All 73 copies contributed, giving 98.6% dominance. Expanded source ambiguity strings remained recorded separately. This assignment was based on the supplied confirmation, rather than candidate order or an assumption that differing reference sequences were identical.

All 52 HLA-B*57:03:01 copies were retained using the available reference sequences: 27 in the group represented by :10, including the compatible shorter :02 reference, and 25 in the group represented by :01. Dominance was 51.9%. Missing terminal reference sequence was left unknown. All 245 HLA-B*51:01:01 copies were assignable; the dominant group contained 231 copies (94.3%). These cases illustrate why reference completeness was distinguished from the sequencing evidence supporting a sample call.

A dagger (†) marks a display representative only when its sequence group contains two or more distinct allele labels in the listed membership. Representatives of single-member groups are unmarked. This convention applies to the main text, main tables and Supplementary Table 1. The symbol is not part of HLA nomenclature; its absence does not establish sequence resolution outside the typing interval. Literal source candidate strings, complete group-member lists, canonical reference accessions and literature-reported allele names remain unmarked. The displayed status “Unresolved within-interval typing (read coverage)” denotes uncertainty in the extended sample assignment: alternative candidates have known differences within the typing interval, but weak or missing read coverage at discriminating positions does not support a single group. This read-coverage explanation was supplied by the study authors; individual read depths and sites were not independently re-evaluated for this analysis. Missing reference sequence is a separate issue and does not establish missing sample reads. The internal category “Conflicting candidate sequences in typed interval” is retained unchanged in machine-readable files for traceability.

All 14,916 class I source-copy records remained in the resource. Of these, 14,716 contributed to group fractions: 13,990 using complete references, 653 using shorter references, and 73 using the confirmed original HLA-B*40 groups. A further 184 allele-copy records retained unresolved within-interval typing ambiguities and 16 records lacked an available genomic reference sequence. Their primary HLA calls remained in the resource, but no single reference-derived group was assigned. No extended call was missing. No separately reported incomplete-reference possible candidates were supplied in the authoritative call export; these annotations were not inferred from earlier exports.

Copies whose ambiguity candidates spanned different known within-interval sequence groups were excluded from group-fraction denominators rather than assigned to the first candidate. For example, eight B*07:02:01 copies had expanded candidate lists containing B*07:02:01:01 and B*07:02:01:10 and remained unassigned, whereas 260 copies supported the B*07:02:01:01† group. Its conditional dominant fraction was therefore 260/260 (100%), with 260/268 (97.0%) of source copies assessed and descriptive dominance bounds of 97.0–100%. The unresolved copies retained their primary three-field B*07:02:01 calls for the other HLA analyses. Candidates differing only outside the typing interval could map to the same group and were not excluded solely because the source contained an ambiguity.

Dominant-group fractions used assigned copies as the denominator. Supplementary Table 1 reports assigned and available copies for every lineage and super-population. Descriptive lower bounds used the leading assigned group count divided by all available extended copies; upper bounds allowed every unresolved copy to join that group. No unresolved identity was imputed. Common lineages had pooled primary frequency ≥1%, and population cells with fewer than five assigned copies were masked. These summaries describe within-assay sequence groups, rather than validating every full-genomic fourth-field distinction.

Eligibility for the common-lineage three-field to fourth-field summaries was determined from pooled primary HLA frequency before excluding unresolved extended copies. The threshold was ≥1% at each locus, equivalent to at least 50 of 4,972 allele copies in 2,486 individuals. It selected 24 HLA-A, 30 HLA-B and 24 HLA-C lineages. A three-field lineage observed once therefore did not enter these common-lineage summaries. No separate minimum count was applied to a fourth-field sequence group within an eligible lineage: singleton groups were retained and contributed to dominant fractions and subtype counts. This distinction avoids suppressing rare subdivisions of a common lineage. The Analysis eligibility sheet in Supplementary Table 1 lists all reported lineages, pooled counts and frequencies, assignment denominators and inclusion flags.

For each eligible lineage, all assigned copies were summarized in the pooled panel and in each super-population. Supplementary tables retain sparse strata with their counts. Population-figure cells with fewer than five assigned copies were masked; mean pairwise Jensen–Shannon divergence used only super-populations with at least three assigned copies for that lineage. These are population-stratum thresholds, not filters on individual fourth-field groups. The predictable (≥95%), mostly predictable (≥80% and <95%) and uncertain (<80%) categories use unrounded conditional dominant fractions. Unresolved identities are not imputed, and no individual is removed from the cohort solely because one extended group is unresolved.

### S2 Annotation of the two leading fourth field representatives

Within each of the eleven globally uncertain lineages, sequence groups were ranked by pooled assigned-copy count. The first and second representatives were taken from exactly the same ranked records as Table 1B. Table 1C was reconstructed from these current pairs, including HLA-B*57:03:01:10 versus HLA-B*57:03:01:01, rather than retaining a previously selected named pair. The original Table 1C annotations were retained in the archive for provenance.

For each pair, known aligned differences were identified only where both references provided sequence. Unknown positions were not called substitutions or deletions. Internal aligned gaps bracketed by known sequence were treated as indels, with adjacent gap/base differences counted as one indel tract. Only differences within the typing interval appear in Table 1C; the complete known-reference contrast records in Supplementary Table 1 include six additional out-of-interval differences, identified by an explicit scope flag.

Positions are one-based from the first base of the canonical genomic records HLA-A*01:01:01:01, HLA-B*07:02:01:01 and HLA-C*01:02:01:01, not ATG-relative nucleotide numbering. Paired bases are written from the first displayed representative to the second, and indels denote the aligned tract without claiming HGVS normalization. Canonical feature annotations mapped each difference to an intron or untranslated region. The ungapped available genomic sequences of all 22 representatives were independently matched to their official allele-specific genomic records, and every known within-interval differing base was checked against the corresponding allele-specific feature annotation.

The eleven within-interval contrasts contained 14 substitutions and one 11-bp indel. Five contrasts involved introns alone, four the 3′ UTR alone, and two both. The A*03:01:01 pair mapped to intron 6, while the A*30:02:01 and A*68:01:02 pairs mapped to intron 7. The B*18:01:01 pair involved introns 3 and 5 and the 3′ UTR; the B*57:03:01 pair involved intron 5 and the 3′ UTR. These mappings replace the earlier location annotations. Neither location nor a fourth-field label was taken to demonstrate a functional effect.

### S3 Biological interpretation of typing coverage

A transcribed 5′ UTR extends from a transcription start site to the ATG start codon; upstream genomic sequence can also include promoter elements. A 3′ UTR extends from the stop codon to the transcript cleavage and polyadenylation site, which can vary between isoforms. Neither a database sequence edge nor an assay endpoint is a universal biological UTR boundary. The present gene-body and proximal-flank coverage supports typing within those intervals, while complete regulatory characterization or particular fourth-field distinctions can require additional sequence.

The proximal HLA-C 3′ interval includes the region of the documented miR-148a interaction (Kulkarni et al., 2011). More distant sites need not be covered: the functional HLA-C Oct1 promoter variant approximately 800 bp upstream of a transcription start site provides an upstream example (Vince et al., 2016). Alternative polyadenylation in HLA-A similarly motivates defining downstream coverage against the transcript endpoints relevant to the biological question (Kulkarni et al., 2017). These examples provide a rationale for extending an assay when needed; they do not establish functional effects for the variants in this study. A named fourth-field distinction requires coverage of its candidate-discriminating positions regardless of their known function.

### S4 Prediction of MICA and MICB from HLA

MICA and MICB genotypes were predicted at up to two reported numeric fields in 2,393 individuals with complete HLA-B/C and MIC calls, retaining expression suffixes. MIC third-field intronic extensions were excluded; original labels remain in the released MIC source-annotation data. Shorter labels (129 MICA and 23 MICB copies) and six MICA copies with retained provisional tags were not completed or renamed. Thus, outcomes evaluate recorded operational MIC genotypes rather than uniformly complete reference identities. MIC prediction used the primary reported two- or three-field HLA types; fourth-field covered-sequence grouping did not alter those inputs. Extended ambiguities were not expanded into separate observed copies. The two copies were unphased; targets were unordered genotypes rather than chromosome-specific assignments. HLA inputs use the reported assignments interpreted against IPD-IMGT/HLA 3.66.0 (October 2026). HLA variant and coverage annotations are supplied with their original source labels alongside the analytical calls in Supplementary Table 7.

B~C~MICA~MICB haplotype frequencies were estimated by EM, which integrates uncertainty in multilocus chromosome phase (Excoffier and Slatkin, 1995). All genotype-compatible unordered haplotype pairs were considered, with random pairing of haplotypes assumed within each training reference. Three initializations were used: the product of training marginal allele frequencies, uniform haplotype frequencies, and a seeded Dirichlet initialization. The converged solution with the greatest training likelihood was retained. Estimation was limited to 3,000 iterations; convergence required both a change in log likelihood per training individual below 0.00000001 and a maximum frequency change below 0.000001. All 35 cross-validation fits converged. Rare alleles were retained, and no frequency-filtered full-cohort haplotype export entered the prediction analysis.

Five folds were constructed by shuffling individuals within each of the 25 sub-populations and distributing them across folds while balancing super-population counts, using seed 20261012. Five folds allowed representation of IBS, which had seven paired individuals, in every test fold. The same folds were used for both MIC genes and four configurations: two-field HLA-B/C with pooled training frequencies; two-field HLA-B/C with frequencies estimated within the known super-population; three-field HLA-B/C with pooled frequencies; and two-field HLA-B alone with pooled frequencies. For the last configuration, pooled four-locus haplotypes were marginalized over HLA-C before conditioning on HLA-B. The population-specific model requires a known super-population label and is therefore an HLA-plus-population predictor. All frequency estimation used training individuals only. No held-out MIC call entered the predictor.

For each held-out HLA genotype, posterior probabilities were summed across compatible haplotype pairs and phase configurations to obtain the marginal distribution of each MIC genotype; the genotype with the largest marginal probability was called. Numerical ties were resolved by allele-label order. Unsupported population-specific inputs used the pooled training conditional model. Remaining unsupported inputs used the empirical pooled training MIC genotype prior. All such calls remained in overall accuracy, and fallback use was reported. The majority-genotype baseline was derived separately within each training fold, using pooled training samples for pooled configurations and the corresponding training super-population for the population-specific configuration.

Exact unordered genotype accuracy was the primary outcome. Unordered allele-copy concordance used the best matching of the two predicted and observed allele copies. Per-allele sensitivity measured recovery of observed copies, and positive predictive value measured the fraction of predicted copies that matched the observed genotype. Macro-averaged allele sensitivity gave equal weight to alleles, complementing overall accuracy in the presence of uneven frequencies. Fold-specific per-allele reporting required at least ten training copies; rare and unseen held-out alleles remained in all overall outcomes. Reporting eligibility and copy counts were supplied separately so that this threshold did not conceal rare-allele errors.

Accuracy among selected calls and call rate were reported together at fixed posterior thresholds of 0.50, 0.60, 0.70, 0.80, 0.90, 0.95 and 0.99. Calibration compared mean posterior probabilities with observed correctness in ten fixed probability bins. Wilson intervals described uncertainty in held-out accuracy proportions (Wilson, 1927), and 2,000 paired bootstrap resamples within sub-population strata summarized accuracy differences from the majority baseline (Efron, 1979). These intervals condition on the fitted out-of-fold predictions and do not include uncertainty from re-estimating reference panels. Full-panel references were fitted only after cross-validation for release; they were not used to obtain the held-out results.

### S5 Cohort weighted fourth field information

The separate cohort-weighted analysis retained every source-reported lineage label, including singleton lineages and singleton assigned groups, without the ≥1% common-lineage filter and used the frozen cohort of 2,486 individuals. At each locus, the most frequent assigned sequence group within a reported lineage defined its modal baseline. Pooled modal groups were estimated over the full cohort; ancestry-specific modes were estimated within each super-population and applied to its own individuals. All modal groups were estimated in the same data being summarized, so these are descriptive counts rather than cross-validated predictions. Ties selected the numerically lowest representative. The complete modal mappings and ties are released in data/results/classI_resolution/weighted_modal_baselines.csv.

We counted non-dominant assigned copies directly and divided by all assigned copies in scope. This equals a frequency-weighted sum of within-lineage residual fractions when lineage weights use the assigned-copy distribution. Bounds using all source copies leave unresolved group identities unknown. Sensitivities retain only common lineages or explicitly three-field primary labels, without completing shorter labels or truncating four-field labels. Among 14,916 primary class I copy labels, 14,816 are explicitly three-field, 96 shorter, and four retain a four-field label.

For individual summaries, the observed HLA-A, -B and -C copies were counted directly; no independence assumption or phase inference was used. An individual with at least one known non-dominant copy was positive even if another copy remained unresolved. Individuals without a known non-dominant copy but with an unresolved copy had indeterminate event status. Lower and upper bounds excluded or included these remaining indeterminate individuals as events. Per-1,000 values rescale these observed bounds within this panel; they are not estimates of a clinical outcome or a prospective new cohort.

All modal-tie combinations were evaluated for the all-lineage individual counts. Four pooled tie combinations all gave 981 known non-dominant individuals; unresolved-event upper counts varied from 1,102 to 1,103. Sixty-four ancestry-specific tie combinations gave 736–749 known non-dominant individuals and unresolved-event upper counts of 877–887. The prespecified numerical convention gives 741 known events and an upper count of 881 for the ancestry-specific analysis. Complete group/copy summaries, individual denominators and tie sensitivities are in Supplementary Table 1. These measures describe additional sequence discrimination within assay coverage; no functional or clinical benefit was tested.
