## Supplementary figures and images for "High-resolution HLA and MICA/MICB typing of the 1000 Genomes panel: a resource for precision medicine"

### SupplementaryFigure1_Typing_workflow.png

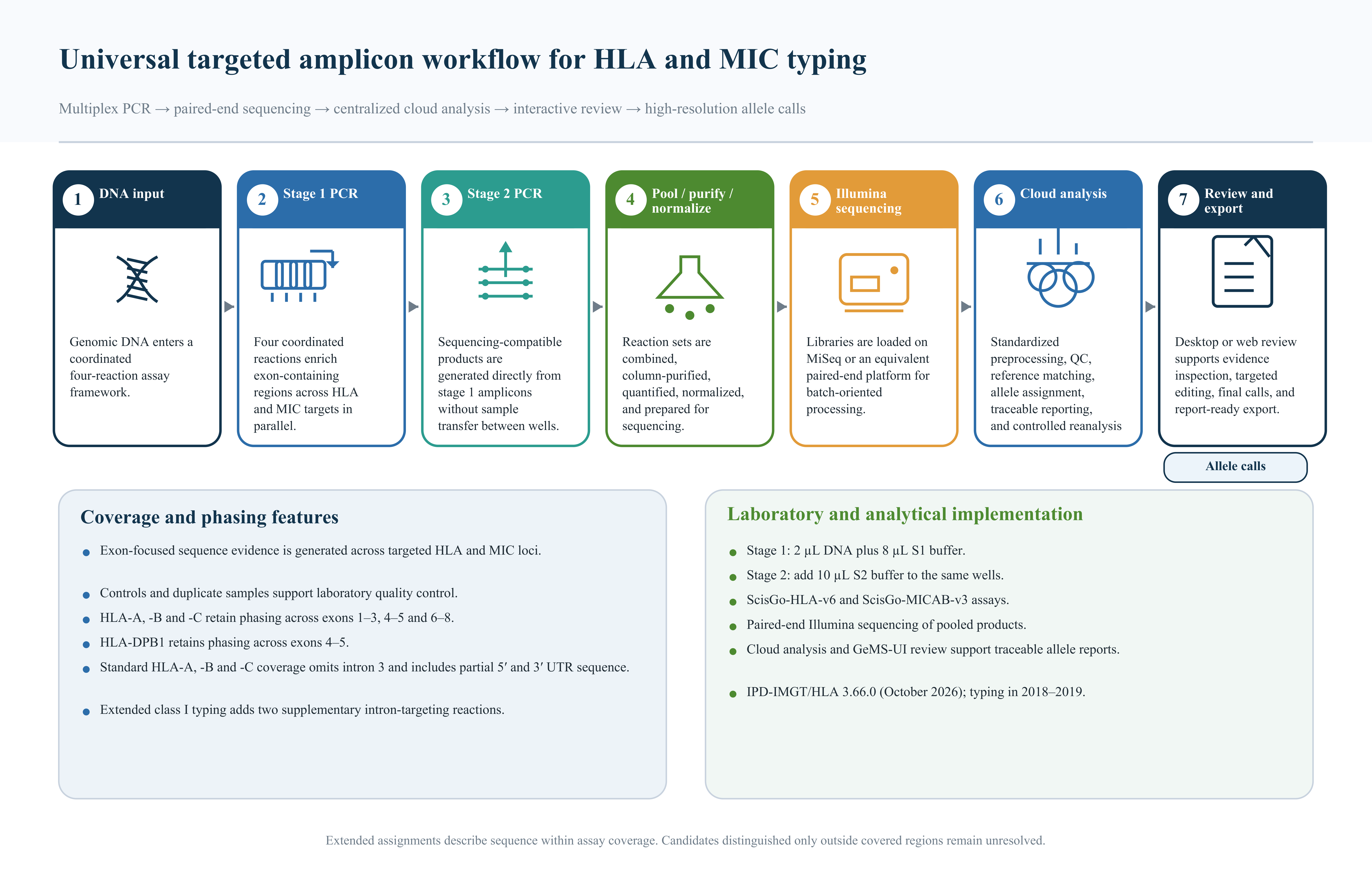

### SupplementaryFigure2_Extended_classII_haplotypes.pdf

**A Extended class II**

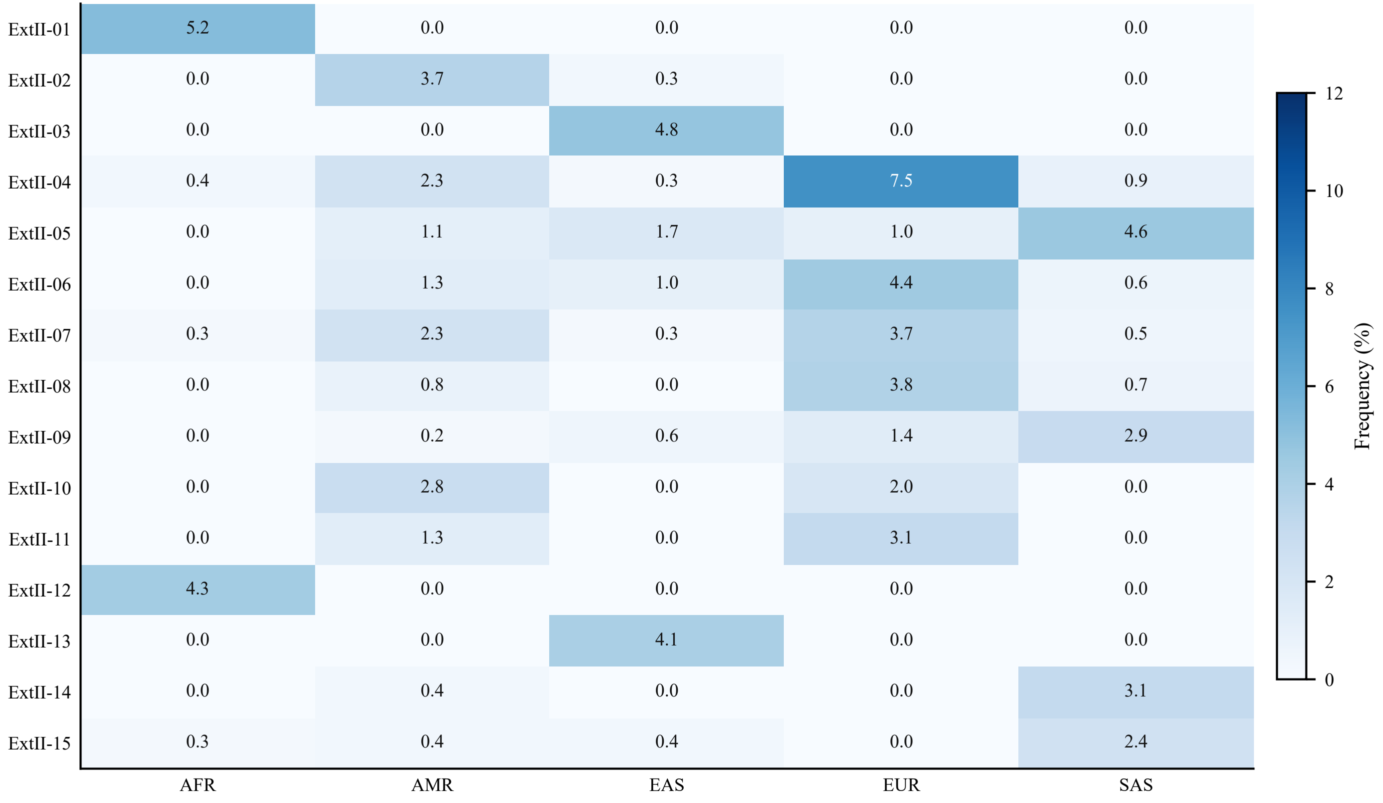

### SupplementaryFigure2_Extended_classII_haplotypes.png

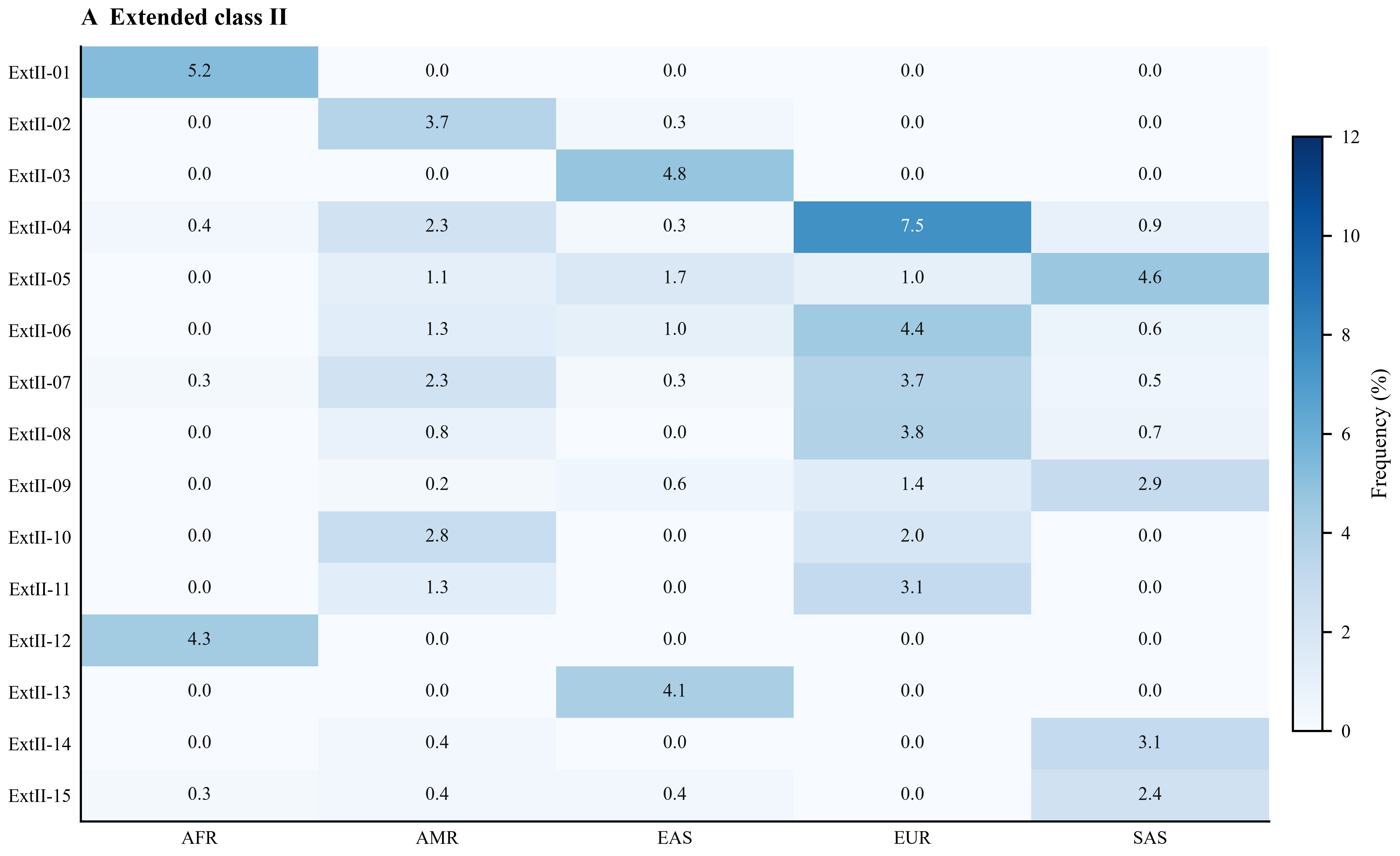

### SupplementaryFigure3_Combined_HLA_MIC_haplotypes.pdf

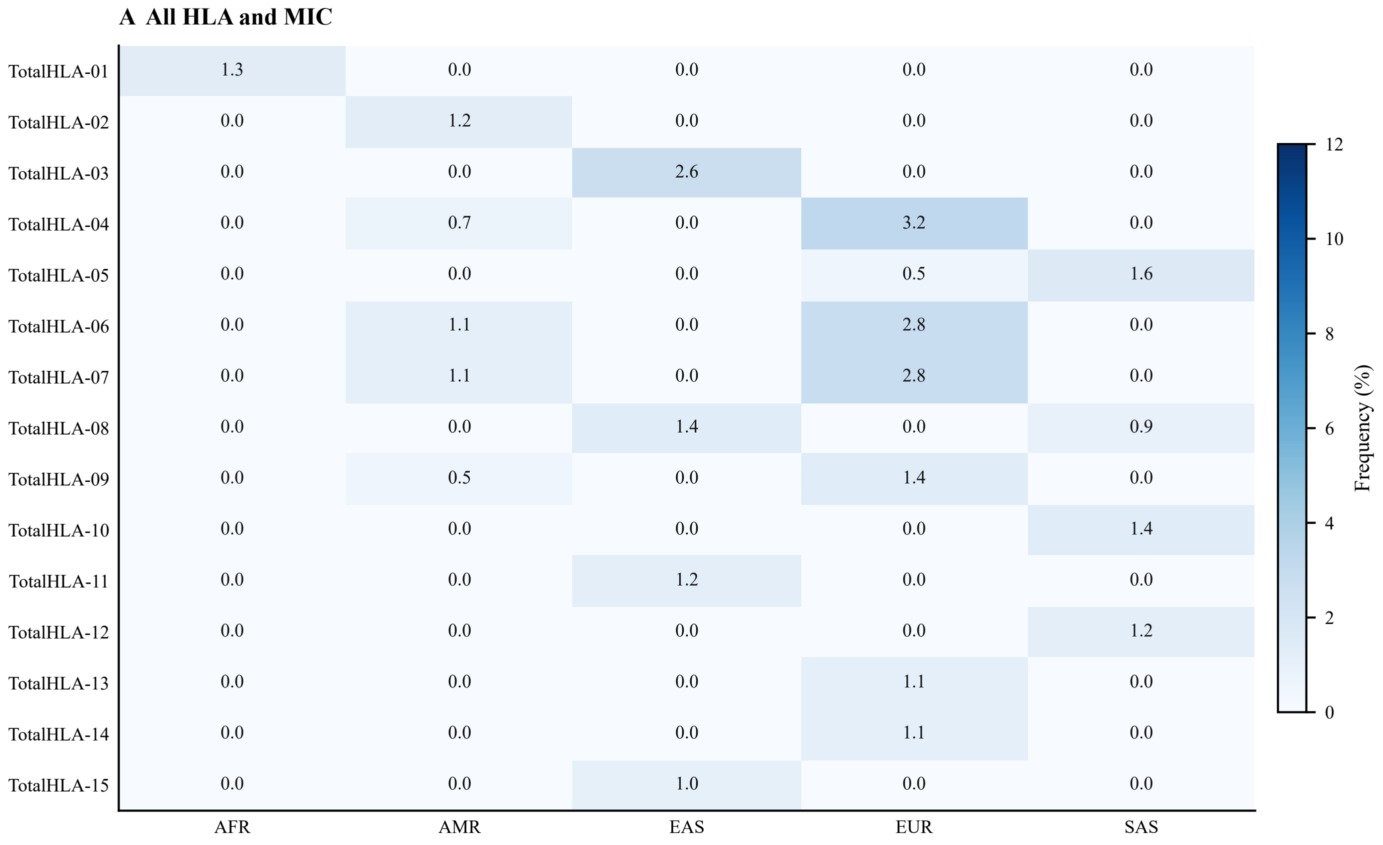

### SupplementaryFigure3_Combined_HLA_MIC_haplotypes.png

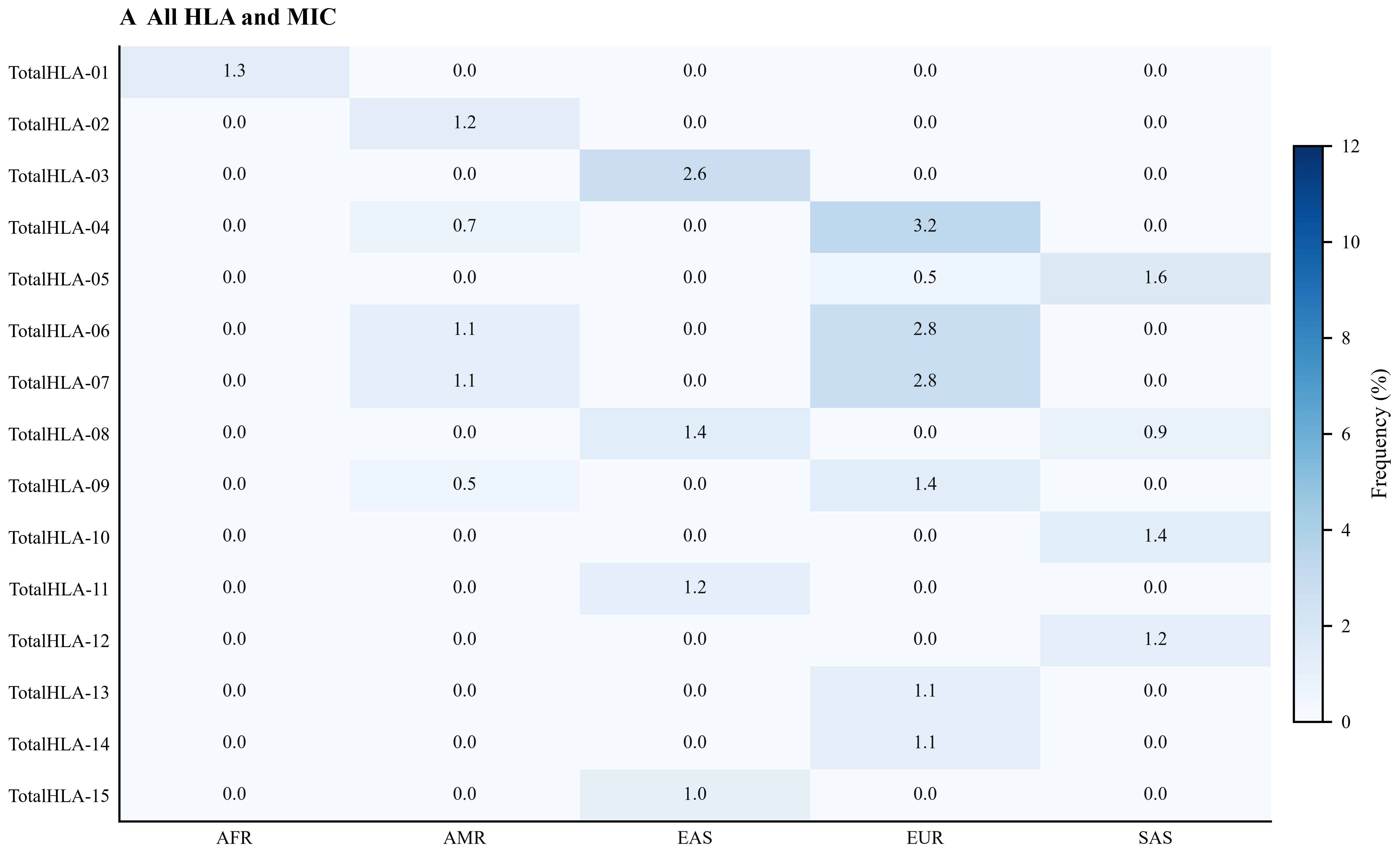

### SupplementaryFigure4_MIC_per_allele.pdf

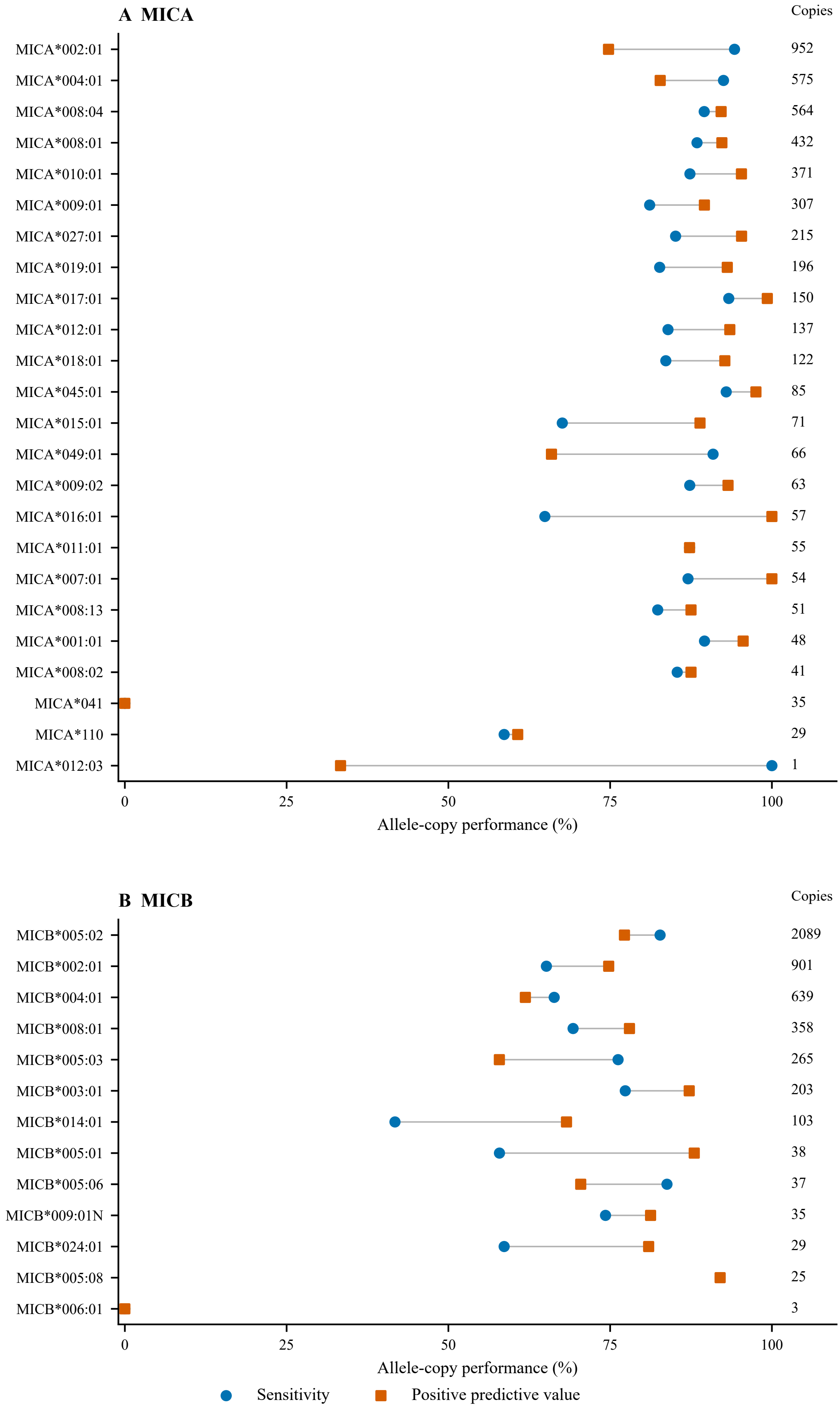

### SupplementaryFigure4_MIC_per_allele.png

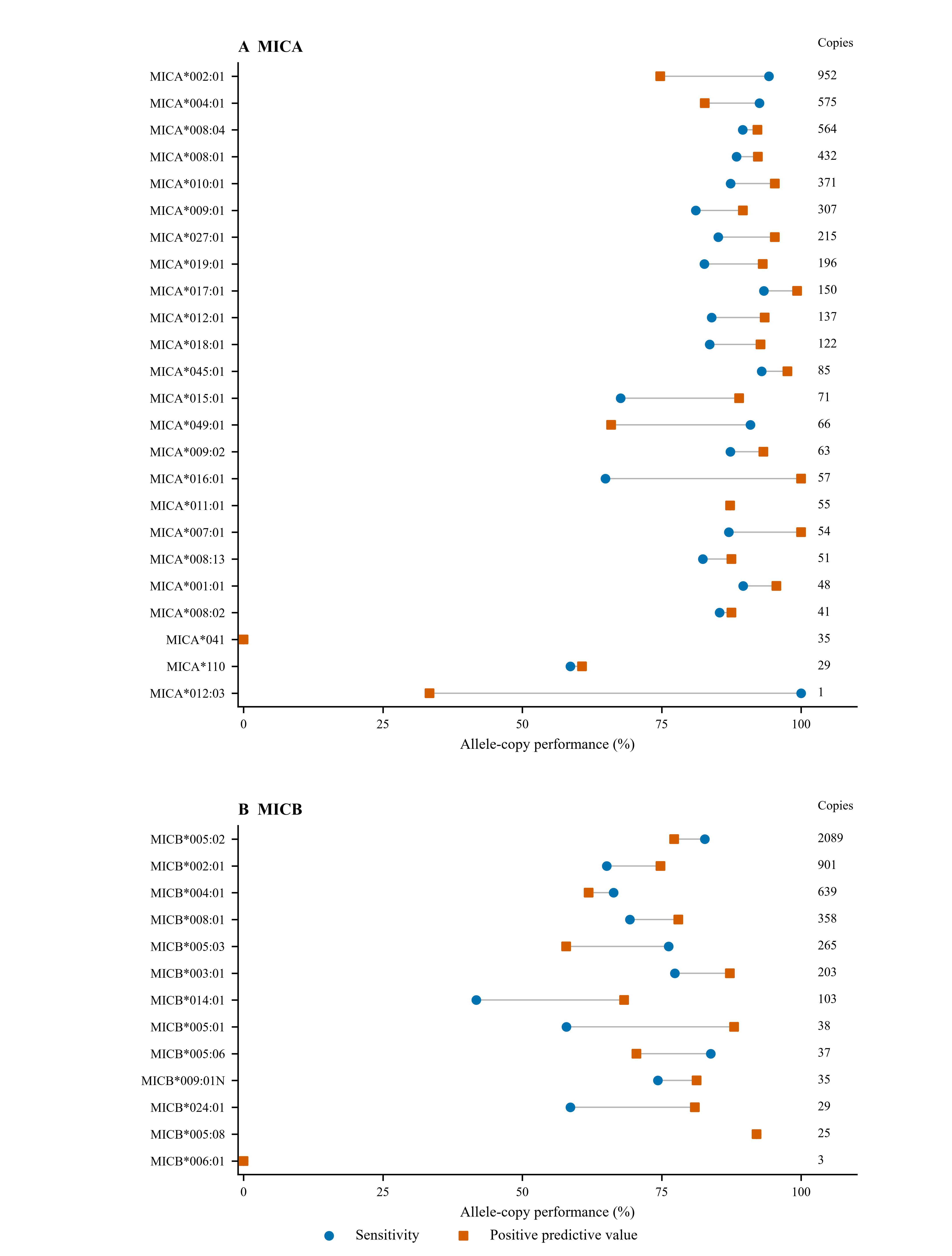

### SupplementaryFigure5_MIC_confidence.pdf

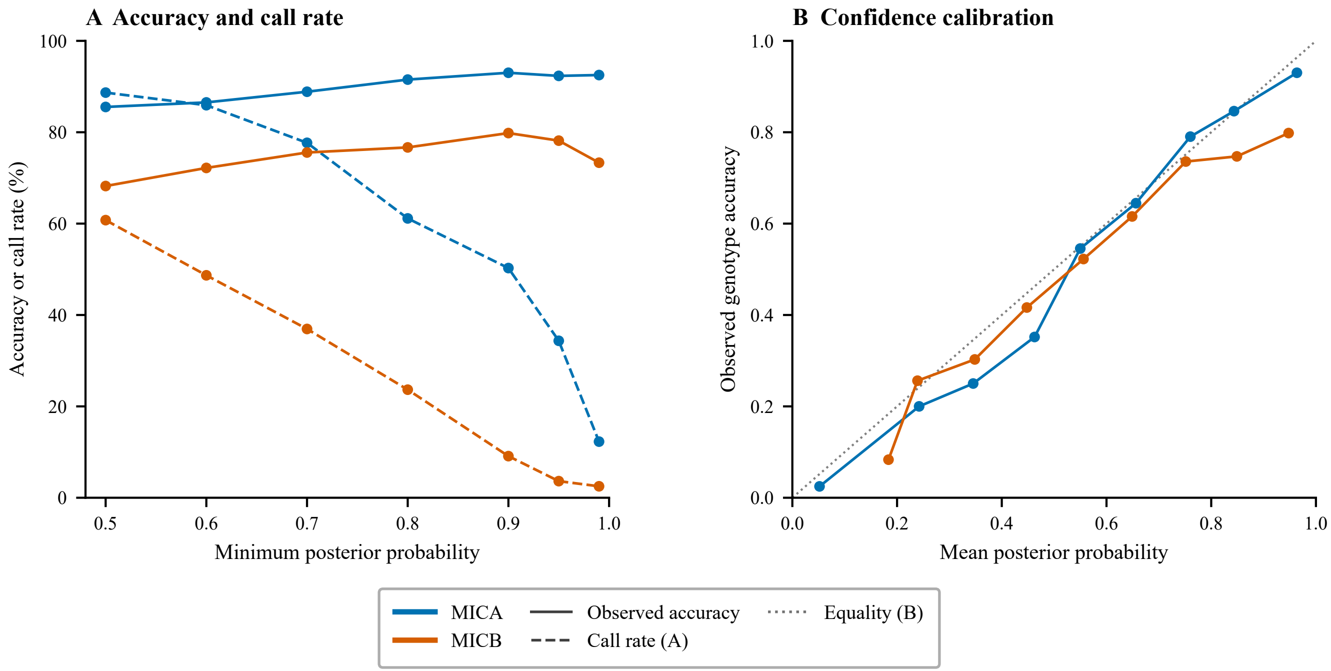

### SupplementaryFigure5_MIC_confidence.png

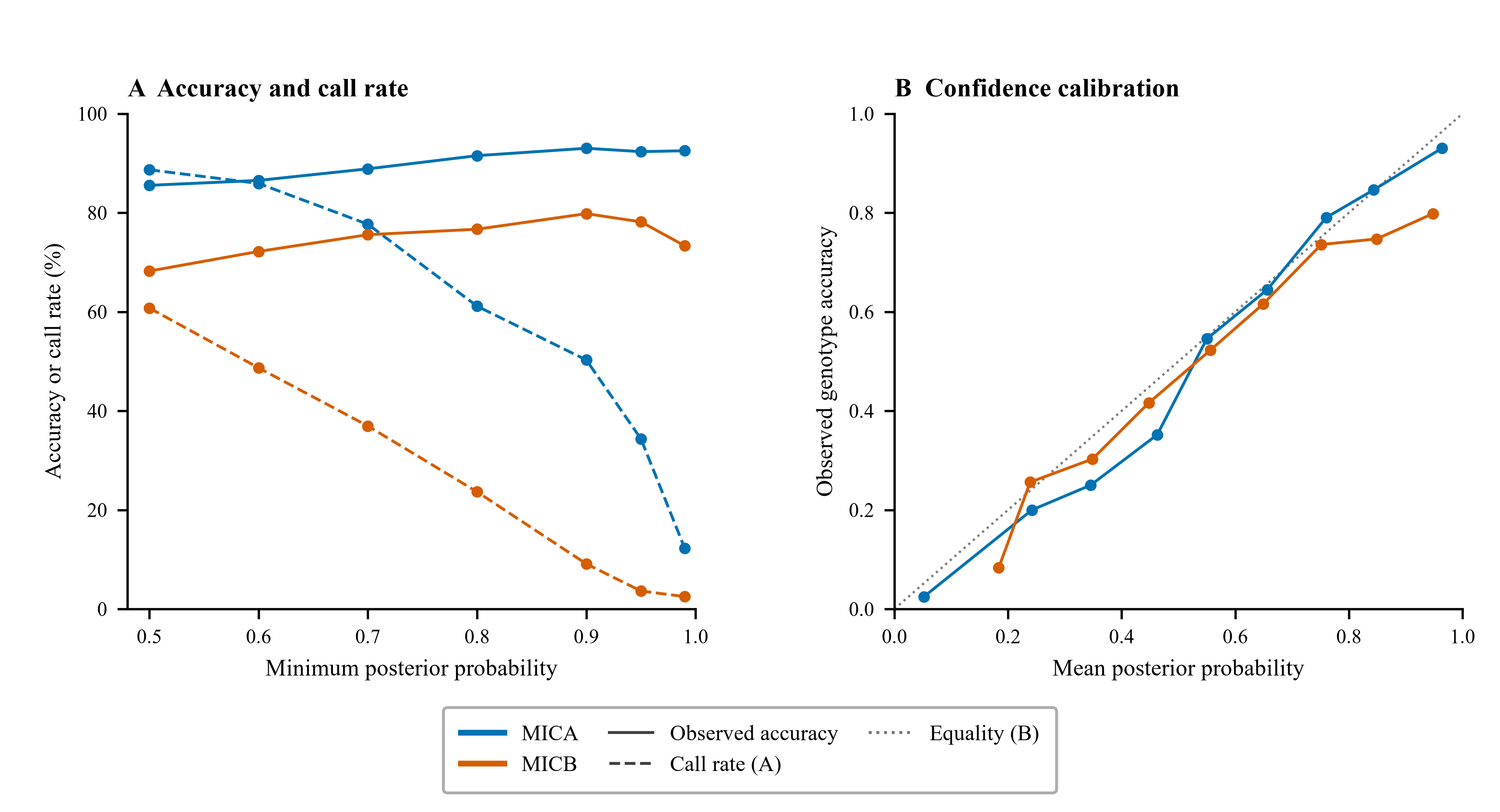

### SupplementaryFigure6_Weighted_resolution.png

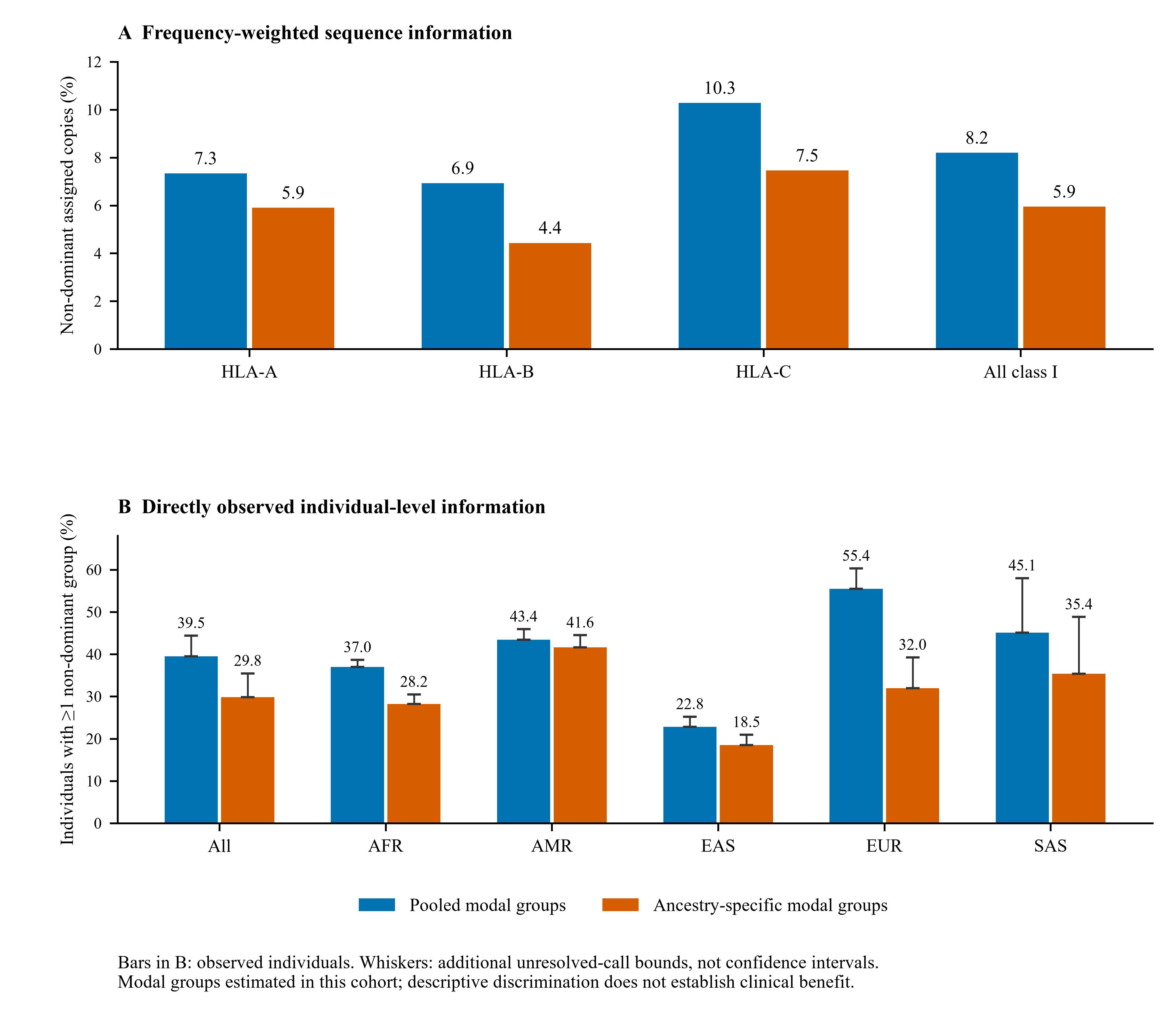
